# Reconciling Membrane Protein Simulations with Experimental DEER Spectroscopy Data

**DOI:** 10.1101/2020.12.19.140186

**Authors:** Shriyaa Mittal, Diwakar Shukla

## Abstract

Spectroscopy experiments are crucial to study membrane proteins for which traditional structure determination methods still prove challenging. Double electron-electron resonance (DEER) spectroscopy experiments provide protein residue-pair distance distributions that are indicative of their conformational heterogeneity. Atomistic molecular dynamics (MD) simulations are another tool that have proved vital to study the structural dynamics of membrane proteins such as to identify inward-open, occluded, and outward-open conformations of transporter membrane proteins, among other partially open or closed states of the protein. Yet, studies have reported that there is no direct consensus between distributional data from DEER experiments and MD simulations, which has challenged validation of structures obtained from long-timescale simulations and using simulations to design experiments. Current coping strategies for comparisons rely on heuristics, such as mapping nearest matching peaks between two ensembles or biased simulations. Here we examine the differences in residue-pair distance distributions arising due to choice of membrane around the protein and covalent modification of a pair of residues to nitroxide spin labels in DEER experiments. Through comparing MD simulations of two proteins, PepT_So_ and LeuT - both of which have been characterized using DEER experiments previously - we show that the proteins’ dynamics are similar despite the choice of the detergent micelle as a membrane mimetic in DEER experiments. On the other hand, covalently modified residues show slight local differences in their dynamics and a huge divergence when the spin labels’ anointed oxygen atom pair distances are measured rather than protein backbone distances. Given the computational expense associated with pairwise MTSSL labeled MD simulations, we examine the use of biased simulations to explore the conformational dynamics of the spin labels only to reveal that such simulations alter the underlying protein dynamics. Our study identifies the main cause for the mismatch between DEER experiments and MD simulations and will accelerate developing potential mitigation strategies to improve simulation observables match with DEER spectroscopy experiments.

## Introduction

Double electron-electron resonance (DEER) spectroscopy has made incredible progress in the study of biomolecules such as cytoplasmic and membrane proteins and nucleic acids (*1*, *2*), including experiments *in vitro* and *in vivo* (*3*–*5*). In DEER experiments, a spin probe is covalently attached to two residues on the biomolecules. Distances between these two spin probes can be determined by measuring the dipolar coupling between an electron pair, one unpaired electron on each of the spin probes. The interaction between electrons is measured in the time domain and then mathematically transformed to distance distributions. Methodological developments have made it possible to obtain distance distributions up to 10 nm in cytoplasmic proteins and 8 nm in membrane proteins (*1*, *6*–*8*), up to 16 nm with sparse spin-labeling that can avoid the deleterious impact of multiple spin labels in close proximity (*9*).

DEER spectroscopy experiments are key for structural insights into membrane proteins for which structure determination methods such as X-ray crystallography and NMR have proved challenging. Given the advance in computational resources, there are numerous extensive molecular dynamics (MD) simulation studies of membrane proteins including GPCRs, transporters, ion channels, integrins, and transmembrane receptor kinases. The observable from DEER experiments, residue-pair distance distributions can be directly compared to dynamics information from MD simulations in order to characterize the structural consequences of the obtained distance distributions. Yet, there is often no direct consensus between distributional data from DEER experiments and MD simulations, which has challenged the validation of structures obtained from long-timescale simulations. Several methods have been introduced to reconcile experimentally characterized distance distributions with simulations such as restrained ensemble MD (reMD) (*10*, *11*) and ensemble-biased metadynamics (EBMetaD) (*12*) simulations, both methods employ the experimentally obtained distance distribution to bias a simulation ensemble. Another method to syncretize unbiased MD simulations with experiments is labeling a residue with a spin probe whose conformational orientations are sampled using a spin probe rotamer library (*13*, *14*). This method is independent of any experiment data bias and relatively computationally inexpensive since no additional simulations are required, but is unable to consider the protein’s conformational dynamics.

Typically, we observe mismatches in terms of relative peak heights when there are multiple peaks in the distance distributions, peak positions, and lower and higher extremes of the distance values. Commonly we observe that experimental distributions exhibit larger distance values, which are not sampled in any of the MD simulation ensembles. These differences can be visualized in Figure S1A where we compare distance distributions from our previous simulations on a peptide transporter protein with experimental DEER distributional data. Most potential for mismatch between experiments and simulation distance distributions stems from differences in experimental conditions and standard simulation protocols. Since membrane proteins are embedded in lipid bilayers in physiological conditions, simulations are typically performed in lipid bilayers. These lipid bilayers can be homogeneous or heterogeneous with different types of lipid molecules (*15*). Bilayer mimetics such as nanodisc (*16*), lipodisq nanoparticles (*17*), bicelle (*18*), liposome (*19*), micelle (*20*) are more amenable to biophysical experiments and have been used for DEER spectroscopy studies of membrane proteins. Specifically, detergent micelles are most commonly used and a widely used detergent is n-Dodecyl-*β*-D-Maltoside (BDDM).

Another significant basis for a mismatch between observed peaks in experiments and simulations is the use of spin probes in DEER experiments, which is absent in wild-type protein simulations. Using site-directed spin labeling (SDSL), two nitroxide spin labels are attached to two cysteine mutated residues. These spin labels can be of different types such as 1-oxyl-2,2,5,5-tetramethyl-pyrroline-3-methyl)methanethiosulfonate (MTSSL), iodoacetamide-PROXYL, unnatural amino acids p-acetyl-l-phenylalanine and 2,2,6,6-tetramethylpiperidine-1-oxyl-4-amino-4-carboxylic acid, and a spin-labeled lysine. DEER experimental measurements among two spin labels are a proxy to explain the protein’s residue-pair distances. Relying on cysteine modifications and the addition of flexible spin probe molecules pose a possibility of modifying the observed protein’s dynamics from DEER experiments. For example, the MTSSL spin probe has five linker dihedrals attributing large rotational flexibility to the protein residue (*14*). Recently metal cations such as Gd^3+^, Cu^2+^ and Mn^2+^ based spin labels that are more rigid have been used (*21*–*23*) but their applications in the study of membrane proteins are limited (*24*).

Based on the above discussed modifications in DEER experiments as compared to physiological conditions, we propose five potential impacts on a protein, its dynamics, and hence the observed DEER experimental observables. Since DEER experiments are typically performed with proteins embedded in bilayer mimetics, such as detergent micelles rather than lipid bilayers, membrane diffusion, packing flexibility and interactions can (1) allow for shifts in DEER distributions and peaks and (2) alter the secondary structure and accessibility of various helices and loops in the protein. Previous studies that draw comparisons between micelle and bilayer environments on membrane proteins have been limited to either small peptides such as single transmembrane helices or are based on ns-timescale simulations that do not provide a realistic picture of a protein’s conformational dynamics. (3) Since DEER measurements require a covalent modification on at least two sites of the protein, we evaluate whether this modulates the underlying free energy landscape of the protein by biasing it to adopt only a subset of the available conformations. In addition to modulating the protein overall dynamics, we examine the impact that the MTSSL probes have locally on the modified residues, their neighboring residues, and their structural properties. (4) We also examine how accurately do the distance distributions obtained from the dipolar coupling of MTSSL spin nitroxide probes provide an approximation of protein dynamics where there is no MTSSL probe. (5) Multiple flexible bonds of nitroxide spin probes (*14*) such as MTSSL spin probes may have different timescales than those from the wild-type residue which will equilibrate at a different timescale than the protein changing the experimentally observed dipolar couplings. We evaluate these perturbations and their impacts in this work to discern which among these is the main cause for the mismatch between experiments and simulations.

Here, we directly compare the biophysical effect of different experimental and simulation conditions by performing MD simulations in conditions similar to experiments. To evaluate the effect of membrane environment on protein structure and dynamics, we compare long-timescales simulations of two proteins in a BDDM micelle and a more typical lipid bilayer. Specifically, we perform simulations of two proteins, PepT_So_ and LeuT, which are biologically important representative proteins of two different membrane protein families, Major Facilitator Superfamily (MFS) and Neurotransmitter: Sodium Symporter (NSS), respectively. Residue pairs in both protein have been previously characterized using DEER experiments (*8*, *20*, *25*, *26*). LeuT has many three-dimensional structures determined through X-ray crystallography and has been investigated using computational simulations. Recently, two crystal structures of PepT_So_ were resolved (*20*, *27*) and we have examined this protein using MD simulations in our previous work (*28*). We follow our micelle and bilayer simulations by introducing nitroxide spin labels MTSSL on a pair of residues in PepT_So_ to examine the perturbations caused by the probe’s site-specific mutations during DEER spectroscopy experiments. We then perform restrained ensemble molecular dynamics (reMD) simulations to evaluate the spin pair equilibration and its impact on the protein’s conformational landscape and residue-pair distance distributions.

## Methods

### MD Simulations

All simulations were setup up using CHARMM-GUI (*29*–*33*), built with a rectangular box and a minimum water height of 10 Å above and below the membrane. System-specific details are provided below. All simulations were run using NAMD 2.13 MD package (*34*) and the CHARMM 36 force field (*35*–*38*) on the Blue Waters petascale computing facility. We used the NAMD inputs generated by CHARMM-GUI for minimization and equilibration in six consecutive steps followed by production runs in the NPT ensemble. The constant temperature was maintained by employing Langevin dynamics with a damping coefficient of 1 ps^−1^. The Langevin piston method was employed to maintain a constant pressure of 1.0 atm with a piston period of 50 fs. Nonbonded interactions were smoothly switched off at 810 Å and long-range electrostatic interactions were calculated using the particle mesh Ewald (PME) method. For all simulation steps, bond distances involving hydrogen atoms were fixed using the SHAKE algorithm. Minimization was done for 10 000 steps, total equilibration and production run time for individual simulations are noted in Table S1. Production simulations were run at 303.15 K, trajectory parameters were determined every 2 fs, and coordinates were saved every 100 ps. All trajectory analysis was done using MDTraj 1.7 (*39*) except where otherwise noted. Analysis methods and workflows are explained in the supplementary material.

### Determining a micelle size for membrane protein simulations

Previous work on simulating protein-micelle complexes (*40*) posits the use of the number of detergents more than the aggregation number of a detergent-only micelle that is 135-145 for the n-Dodecyl-*β*-D-Maltoside (BDDM) detergent (*41*, *42*). Moreover, the BDDM micelle size was determined to be 72 kDa (*43*); with a 510.621 g/mol molecular weight of a BDDM molecule (*44*) this yields ~141 detergent molecules in the micelle. To test the stability of the protein-micelle complexes, we took a single structure of the PepT_So_ protein from our previous simulations (*28*) and embedded it in 150, 180, and 200 BDDM detergent molecules. The three simulation setups with 145 668, 145 847, and 146 061 atoms respectively comprised of protein, detergents, waters, and 0.15 M KCl ions. Simulation setup with 150 detergent molecules was minimized for 10 000 steps and the other two were minimized for 20 000 steps. We ran each of these simulations for 60 ns each post-equilibration and only the last 50 ns were used for analysis (Table S2) to assess the protein’s structure and dynamics in all three micelle sizes. RMSD of the protein converges to values between 0.28-0.32 nm within 50 ns, and these values are lower when only the transmembrane region of the protein is included (Figure S2A). We do not expect to sample any conformational change in the protein’s structure in such short trajectories.

We then evaluated the extent of sphericity of the micelle, measured by calculating its eccentricity where a perfectly spherical object has eccentricity 0. We find that in all three cases, micelles in our simulations are spherical with an average eccentricity of 0.23 (±0.02) for micelle with 150 and 200 detergents and 0.22 (±0.02) for micelle formed by 180 detergent molecules (Figure S2C). The radii of the micelles do not show much variation, indicating that the micelles do not distort (Figure S2D). As expected, micelles with more detergents have a larger average radius - 4.4 nm, 4.58 nm, and 4.68 nm for 150, 180, and 200 detergent micelles, respectively. Figure S2E shows a radial distribution of distances between BDDM detergent molecule headgroups. Since the distribution is the same for all three micelle sizes, we conclude that detergent packing is similar in all three micelles.

Our preliminary simulations indicated that protein dynamics and shape of detergent micelle do not vary with the number of detergents in the micelle. We chose 150 detergent micelle for the rest of our simulations to keep the system sizes smaller and conserve computational resources. We were also able to confirm that the simulation setup is stable for all three micelle size choices.

### Setting up LeuT simulations in bilayer and detergent micelle

We compiled 28 LeuT crystal structures among which all but two have no mutations in the proteins sequence. Structures 3TT1, an Outward Facing (OF) structure with two mutations, and 3TT3, an Inward Facing (IF) structure with four mutations (*45*), were modeled on the wild-type LeuT sequence using Modeller (*46*) interface in Chimera (*47*). A recent study by Gotfryd et al. reported a LeuT IF-occluded conformation without any mutations (*48*) but the study and the structure were released after the simulations in this work had been performed. Based on these PDBs, we identified 36 unique structural models for the LeuT protein from residue Arg5 to Ala513 (509 residues in all). Most of these 36 structures were missing residues either on EL2, EL3, or EL6, and the size of the largest missing region in any structure was six residues. These missing regions were modeled to yield 72 LeuT structures as a starting point for our simulations. These structures were aligned in VMD (*49*) using orient and a linear algebra Tcl package, La. During setup in CHARMM-GUI, LeuT structures were capped with ACE and CT3 residues. For protein-bilayer complexes, the structures were embedded in a POPE bilayer of 150 lipid molecules equally distributed in the upper and lower leaflet using the Insertion method. For protein-micelle complexes, protein structures were embedded in 150 BDMM detergent molecules. We only added three Cl^−^ ions to neutralize the system. Since we are only interested in the equilibrium conformational changes of *apo* protein, we did not want to introduce Na^+^ ions that are known to play an important role in the transport mechanism of LeuT. Ion binding and substrate transport are coupled and ions can be considered as a substrate.

### Setting up PepT_So_ simulations in bilayer and detergent micelle

We used 42 structures extracted from our previous simulations of PepT_So_ (*28*) from residue Pro8 to Tyr512 (505 residues in all). During setup in CHARMM-GUI, the protein was capped with ACP and CT3 residues. For protein-bilayer complexes, the structures were embedded in a heterogeneous POPE/POPG (3:1 ratio) bilayer of 200 lipid molecules equally distributed in the upper and lower leaflet. For protein-micelle complexes, protein structures were embedded in 150 BDMM detergent molecules. We added 0.15 M NaCl ions in addition to neutralizing the system.

### Setting up PepT_So_ Simulations with MTSSL probes on a single residue pair

The same method as for PepT_So_ in detergent micelle was followed, and residues Asn174 and Ser466 were mutated to MTSSL (1-oxyl-2,2,5,5-tetramethylpyrroline-3-methyl methanethio-sulfonate) (*50*) probes. This corresponds to one of the extracellular residue pairs chosen by Fowler et al. for DEER experiments (*20*). Here, we have used the WYF parameter for cation pi interactions as available in CHARMM-GUI.

Details for system size and simulation time are provided in Table S1. For all simulations described above we examine convergence by randomly sampling 25%, 50%, and 75% of the trajectories and graphing experimental residue-pair distance distributions shown in Figures S3-S7. We choose to look at these residue-pair distance distributions, as a check for convergence, because these will be the main focus of most results in this work. We see that for all systems, error bars are small even with 25% simulation data, and they decrease as we include a larger portion of the data. We conclude that multiple trajectories sample each region of the conformational ensemble and no single trajectory shifts the distance distributions completely.

### Restrained-ensemble molecular dynamics (reMD) simulations for PepT_So_

We used 42 different protein conformations as a starting point for reMD simulations in vacuum which means the protein was not surrounded by lipids, water, or ions. CHARMM-GUI’s default 25 spin-label copies were attached to each labeled protein residue. Experimental distance distributions from Fowler et al. were provided as target histograms (*20*). We also used default values for force constants, bin widths, and Gaussian natural spread (*33*). Simulations were run using a special version of NAMD 2 (*11*, *34*). For system reMD (1 dist), we attached MTSSL probes on residues Asn174 and Ser466, and restrained this distance. For system reMD (2 dist), MTSSL probes were placed on four residues and two distances were restrained, Asn174-Ser466 and Arg201-Glu364. These residue pairs are on the opposite side of the protein. For system reMD (8 dist), MTSSL probes were places on 12 residues, and eight experimentally studied residue pairs were restrained. Details for system size and simulation time are provided in Table S3. We used an integration timestep of 1 fs and saved trajectory coordinates at a frequency of 50 ps. Since reMD simulations are biased simulations, where the distance between the probe molecules is restrained to a targeted distribution using harmonic forces we use these simulations as an opportunity to explore the protein and the MTSSL probe dynamics when the experiment and experiment distance distributions show a perfect match. This is also why reMD simulations in vacuum are efficient and sufficient to sample the spin probe dynamics.

## Results

### Residue-pair distances from proteins in micelle resemble trends in bilayer-embedded proteins

PepT_So_ is a proton-coupled bacterial symporter for which, recently, researchers characterized eight inter-residue distance distributions using DEER (*20*). There are two known crystal structures for this protein found in the bacteria *Shewanella oneidensis*, 2XUT (*27*), and 4UVM (*20*), both in the inward-facing conformation of the protein. PepT_So_ belongs to the Proton-dependent oligopeptide transporter (POT) family and the Major Facilitator Superfamily (MFS) whose members have a wide variety of functions and are found in many different organisms including humans. All MFS transporters share a common structural fold consisting of 12 transmembrane helices (*51*), however, POT family transporters within MFS often have two additional helices. Like most POT family transporters, PepT_So_ has 14 transmembrane helices, divided into N terminal bundle (TMs 1 to 6), C terminal bundle (TMs 7 to 12), and two extra helices A and B between the two domains packed closely with the C terminal bundle.

LeuT, a leucine transporter, has many high-resolution crystal structures and has been extensively characterized using DEER experiments (*8*, *25*, *26*). LeuT belongs to the Neurotransmitter: Sodium Symporter (NSS) family whose other members include Dopamine, noradrenaline, GABA, glycine, and serotonin transporters. LeuT was the first structure resolved using X-ray crystallography from the NSS family. Although many structures have been resolved since then, only one structure is inward-facing as a quadruple mutant (3TT1 (*45*)). The LeuT-fold consists of 12 transmembrane helices, of which TMs 1 to 5 and TMs 6 to 10 are inverted repeats of each other.

PepT_So_ and LeuT are model proteins from two different families of membrane proteins. While LeuT has been studied using computational simulations with both unbiased and biased protocols, there are only a few short-timescale computational studies focused on PepT_So_. Our previous work sampled the conformational dynamics of PepT_So_ using long-timescale 54 *μs* MD simulations and analyzed its equilibrium dynamics using Markov state model based analysis (*28*). These simulations were carried out in a POPC bilayer using the AMBER FF14SB force field. To compare the dynamics of PepT_So_ protein in detergent micelle and bilayer and solely capture the effect of the membrane environment, we replicated our simulations in a POPE/POPG bilayer in CHARMM 36 force field. Simulations from our previous work (*28*) provide a benchmark for sufficient conformational sampling since we were able to sample IF, OC, OF, and multiple other intermediate states (Figure S8). Here, we compare our atomistic molecular dynamics simulations of PepT_So_ and LeuT in BDDM micelles and lipid bilayers.

In Figure S8A,B, we project our PepT_So_ simulation datasets on gating residue pairs, Ser131-Tyr431 on the intracellular side and Arg32-Asp316 on the extracellular distance. We compare the sampled regions with our previous simulations (Figure S8D) to conclude that all physical conformations of the protein have been well sampled. Similarly, in Figure S9 we project our LeuT simulation datasets on one residue pair each on the intracellular and extracellular side of the protein, Arg5-Asp369 and Arg30-Asp404, respectively. These residue pairs are based on gating residues identified in hSERT (*52*). This human serotonin transporter has a typical LeuT-fold and shares 35.5% sequence similarity with LeuT protein. Among the gating residues, Asp404 from LeuT is homologous to Glu493 in hSERT and the other three residues are arginines.

Upon comparing simulated and experimental distance distributions from our micelle and bilayer simulations (Figure S1B,C) we see distance distributions obtained from micelle simulations are no better at matching with experiments. However, by comparing distance distributions in Figure 1, we examine the impact of the choice of membrane on the protein’s dynamics. For PepT_So_ protein, five out of a total of eight distance distributions show a higher median value (middle horizontal line on violin plots in Figure 1A) in micelle as compared to bilayer. For instance, residue pair 174-466 shows a single peak in the distributions for both micelle and bilayer, but the data has a median value of 3.87 nm mean in bilayer whereas this value is 4.00 nm in micelle. On the other hand, two distance distributions for residue pairs 47-330 and 174-401 show lower median values in the micelle than in the bilayer. One distance distribution for residue pair 141-438 shows about the same value 1.6 nm in both micelle and bilayer. In Figure S10, we show that most of the mean or median values lie along the black dotted line, indicating that they are similar for micelle and bilayer. Mean and median values for all inter-helix distances also fall along the dotted line indicating that the differences are minimal.

**Figure 1:**
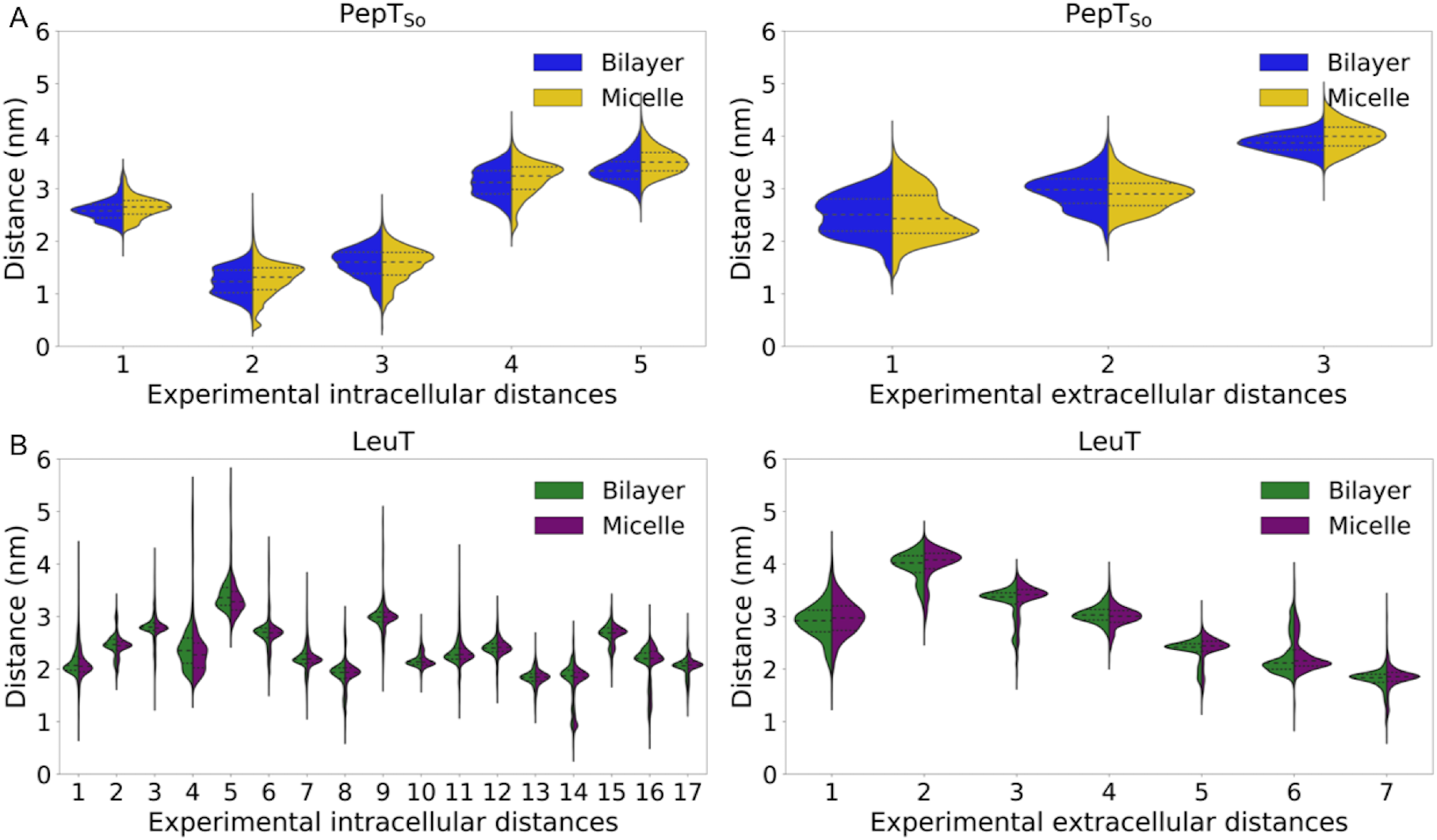
(A) Violin plot shows distance distributions for five intracellular residue-pair distances and three extracellular residue-pair distances measured by Fowler et al. as observed from MD simulations of PepT_So_ protein in micelle (yellow, right) and bilayer (blue, left) (*20*). (B) Violin plot shows distance distributions for 17 intracellular residue-pair distances and seven extracellular residue-pair distances measured by Kazmier et al. as observed from MD simulations of LeuT protein in micelle (purple, right) and bilayer (green, left) (*26*). Distance distributions among *C_α_* atoms and sidechain atoms is shown in Figure S21 and S22.

Based on visual inspection, not only the positions of the peaks but the number of peaks in the distance distributions can differ such as three peaks in bilayer versus two in micelle for residue pairs 141-432 and 141-500. Interestingly, these two distance distributions show new peaks in micelle where little or no data is seen in those regions in bilayer. For LeuT, although the distance distributions differ, the variation is much less (Figures 1B and S10), for example, none of the 24 experimental distance distributions show a peak in bilayer which is not there in micelle or vice versa.

For PepT_So_, five of the experimental residue-pair distance distributions also show slightly broader distributions. For inter-helix distributions (Figure S10) we see that few upper values and lower values lie below the dotted line meaning that the distributions move towards larger values in micelles. Does this mean that micellar environments shift the distributions to larger values? This is unlikely because for LeuT we see values that are both above and below the black dotted line in experimental distances as well as inter-helix distances.

To conclude that the reason for the mismatch between DEER experimental observables and MD simulations distance distributions is due to the use of detergent micelle in MD simulations, distance distributions should have exhibited a consistent behavior such as micellar distributions are always larger, smaller, wider or narrower. However, there are no dramatic or homogeneous shifts in the distance distributions from our simulations.

### Proteins in micelles and bilayers show structural similarity

For both proteins, we measure and show the helicity of transmembrane helices in Figure S11. Values closer to 1 indicate helical nature and decreasing values show loss of helicity. TMs 7 and 10 exhibit a wider range of helicity in PepT_So_ which indicates their dynamic nature. In Selvam et al. we report that one of the extracellular gating residues is on TM7 and one of the intracellular gating residues is on TM10 (*28*). Given that the median of TM7 helicity is 0.76 in both micelle and bilayer, lowest among all other transmembrane helices, none of the helices lose their entire alpha-helical nature. Moreover, broader distributions for TMs 7 and 10 are seen in both micellar and bilayer environments.

TMs 1 and 6 in LeuT show wide helicity ranges in both environments. Readers must note that TM1 here indicates residues of TM1a, the first half of TM1 helix. TM1a is of particular interest in LeuT and other NSS family transporters (*45*, *48*, *53*) because in IF structures this region is away from the bundle as shown in Figure S12C. Low values of TM1 helicity arise from IF trajectories and other trajectories that transition to IF like states. Our simulation ensemble includes two independent trajectories based on the IF structure 3TT3 (*45*). LeuT TM1a dynamics show a significant distinction in OF and IF states in our MD simulation trajectories (Figure S12), TM1a helicity drops to 20-30% in IF trajectory whereas this is 50-80% in OF trajectories. Due to the dynamic nature of this region, it follows that one of the gating residues on both the intracellular and extracellular side of the protein are also positioned on TM1. This distinct behavior of TM1a is also seen in Figure S9A,B where LeuT is open on both extracellular and intracellular sides. Other studies on transporter proteins using extensive MD simulations (*52*, *54*, *55*) have also reported observing this hourglass-like state of the transporter. Terry et al. have reported evidence for this conformation in LeuT which is due to a weaker coupling between the extracellular and intracellular side of LeuT (*56*). We suggest that this weaker coupling allows LeuT to explore a large range of intracellular gating distance while the extracellular side of the protein is also open.

TM regions of PepT_So_ and LeuT show structural similarity in both micelle and bilayer, but could the choice of the membrane milieu affect the intracellular and extracellular flanking regions of our proteins? We compare the distributions for these regions such as the helicity of two short helices in PepT_So_ one on each side. For LeuT, we compare the helical content of the loop regions which connect the TM helices. Figures S13 and S14 show that distance distributions are similar and not impacted by the choice of membrane environment. In this work, we do not consider the molecular-level differences in protein residue interactions with lipids or detergents and differences in membrane curvature that could vary stability of loop regions.

Figure 2 strikingly shows that TM helicity median and mean values lie along the black-dotted line, and in most cases, lower and upper values also don’t deviate much in micelle and bilayer. In general, helicity values or distributions are not different which means that micelles do not impact the structure of the protein.

**Figure 2:**
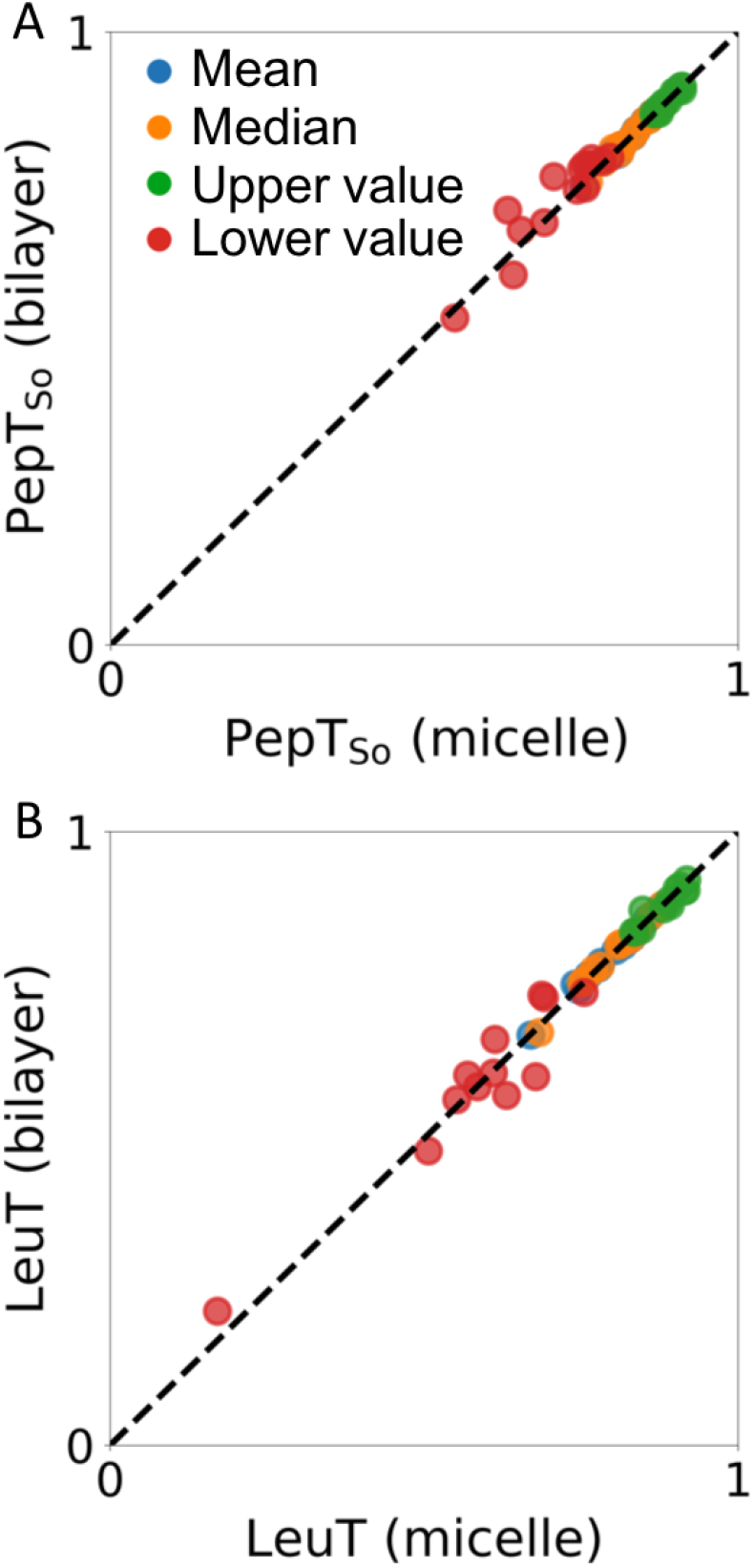
Comparing mean (blue), median (orange), upper value (green), and lower value (red) of alpha-helical content of (A) 14 TM helices in PepT_So_, and (B) 12 TM helices in LeuT. Markers below the black dotted line indicate larger values observed in micelle environment. Markers above the black dotted line indicate larger values observed in bilayer environment. Markers along the black dotted line indicate similar observations in micelle and bilayer simulations.

### Covalent modification due to MTSSL probes cause small local structural perturbations on the protein

We used Kullback-Leibler (KL) divergence to quantify mismatches and differences among the distance distributions. KL Divergence for two distributions *P* and *Q* is 0 if and only if *P* and Q are exactly the same. We calculated KL divergence among distance distributions from MD simulations and micelle and MD simulations in bilayer discussed above. Among eight experimentally characterized distances in PepT_So_, we found that residue pair Asn174-Ser466 has the highest KL divergence value. Hence, we chose this residue pair for further study, specifically to perform simulations with realistic nitroxide DEER labels. We attached an MTSSL DEER probe on Asn174 and Ser466 after mutating them to cystines via CHARMM-GUI these residues and simulated our protein in a BDDM micelle for ~19 μs. Previous studies have compared MD simulations with DEER experimental results with short-timescale MD simulations with explicit spin probes in a variety of biological systems. To our knowledge, this is the first study of the impact of MTSSL spin labels on a membrane protein using long-timescale unbiased simulations.

Figure 3A shows the simulated conformational ensemble projected on the intracellular and extracellular gating residues. Comparing this landscape to those for the PepT_So_ simulations without probes in micelle shows that both ensembles capture all conformational states of the protein. This follows that probe molecules do not seem to interfere with the conformational dynamics of PepT_So_ protein in a way that could hinder its transport function.

**Figure 3:**
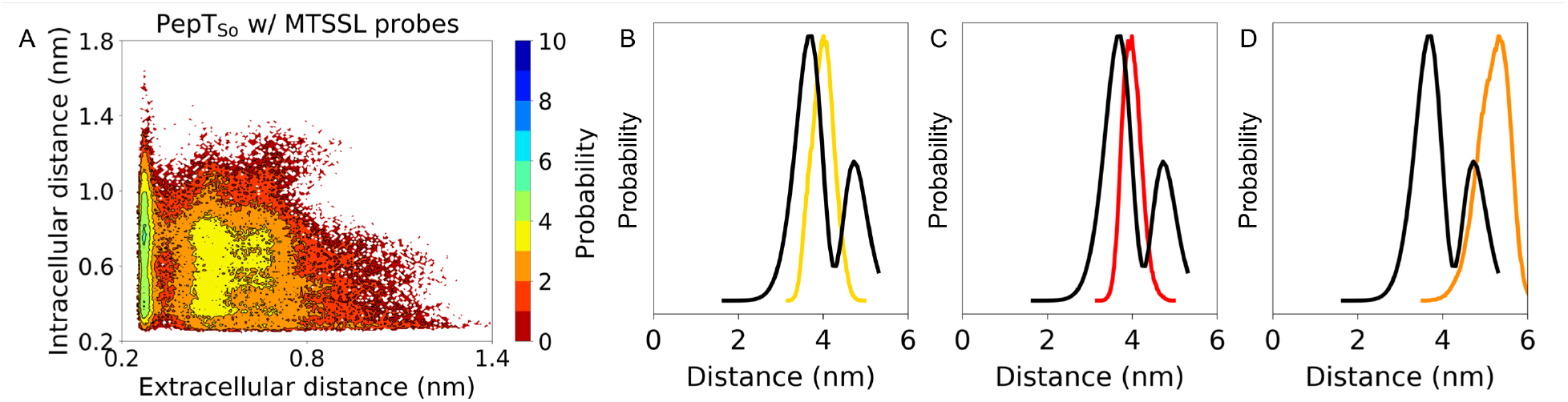
(A) The conformational landscapes of PepT_So_ protein are generated by projecting all simulation data on the chosen extracellular and intracellular side distances measured between Arg32-Asp316 and Ser131-Tyr431, respectively. Conformational landscape for PepT_So_ MD simulations in BDDM micelle with an MTSSL labeled residue pair. (B) Distance distribution for MTSSL labeled residue pair in PepT_So_, 174-466, from simulations in BDDM micelle without probes (yellow), and (C) with probes (red). (D) Distance distribution for MTSSL labeled residue pair in PepT_So_, 174-466, from simulations in BDDM micelle without probes (orange) where distances are measured between ON atoms on MTSSL labels. Black lines show DEER experiment distance distributions.

To understand the local effects of the MTSSL probes on the protein, we calculate Phi & Psi dihedral angles and generate Ramachandran plots for the mutated residues 174 and 466. We see a slightly larger coverage for residue 174 with the MTSSL probe (Figure S15B) as opposed to when it is an Asn residue (Figure S15A), while there is no difference for residue 466. Similarly, when we look at the regions surrounding the labeled residues, specifically two residues both before and after the labeled residues, we see a larger distribution for residue 174 (Figure S15E,F). Hence, we conclude that this mutant created for DEER spectroscopy experiments slightly impacts the local dynamics and secondary structure of the protein, this effect does not alter the overall conformational dynamics of the protein.

We suggest that any alteration seen in transport activity could be due to the kinetic rates of the transport function that would not affect the DEER observations unless functional interactions are mutated. Fowler et al. tested the transport activity of their PepT_So_ double cysteine mutants and 174-466 mutant although decreased activity, did not abolish AlaAla transport entirely (*20*). Kazmier et al. also tested binding of Leu to spin-labeled LeuT pairs and most double mutants retained more than 50% binding affinity as the wild type protein.

### MTSSL probe distances are vastly different when compared to distances from wild-type protein simulations

We examine the impact of a spin-probe labeled residue pair on the resulting distance distributions (Figure S16) by comparing micelle simulations with and without MTSSL probes. Since the probe molecules are on the extracellular side of the PepT_So_ protein, two of the three extracellular side distance distributions do appear slightly perturbed. We observe that the intracellular side distances show no differences. A closer look at the distribution for the residue pair labeled with MTSSL probes shows that the median values, 25%, and 75% quartile values are conserved. Overall, we don’t see any significant changes in the distance distributions for all eight experimental distances as compared to the distance distributions obtained from simulations in BDDM micelle. As expected, if there is no overall difference in the underlying conformational landscape as we discussed above, individual residue-pair distances also would not deviate. Quantitatively, symmetrized divergence values indicated that distance distributions from MD simulations in micelle with and without probes were less divergent as compared to distance distributions from MD simulations in bilayer and micelle.

Most nitroxide spin labels such as MTSSL consist of an unpaired electron delocalized over the N-O bond. Dipolar coupling can be estimated by measuring the distance between two oxygen atoms (referred to as ON atom) from our simulations with two MTSSL labeled residues. Figure S17 shows the ON-ON atom distance distributions as compared to the *C_α_* and sidechain atom distances as observed in these probe simulations. Given the long length of the MTSSL molecule it is expected that ON-ON atom distance distributions are upward shifted with a median value of 5.22 nm. On the other hand, the median value of the closest heavy atom based distance distribution is ~0.9 nm lower than the ON-ON atom distance median.

Comparing these distributions to the experimental DEER distribution, we see single peaks from MD simulations whereas two peaks are seen in the experimental distribution, black lines in Figure 3. From Figures 3B and C, it appears that the MD simulation data capture conformations corresponding to the first peak, but Figure 3D shows that the ON-ON atom distance distribution points to conformations captured corresponding to the second peak with a larger distance value. Evidently, the choice of the atom for distance calculations is imperative and may significantly impact structural inferences drawn from MD simulations. While the ON-ON atom distance distribution changes our view of which peak our data corresponds to, we still do not match the experimental data and nor do we obtain multimodal distributions as seen in experiments.

### Restrained-ensemble MD simulations sample spin probe dynamics, but alter protein dynamics

Our results above elucidate that MTSSL probes modulate the distance distributions obtained from DEER experiments and the experimentally characterized distance distributions are a function of both the protein’s dynamics as well as the probe’s dynamics. MTSSL spin labels are long and flexible molecules and their dynamics have not been examined previously over a long time. We believe that our previous simulations are not sufficient to capture the dynamics of the probes and the proteins together, making unbiased simulations intractable to explore MTSSL probe dynamics. Restrained-ensemble MD (reMD) simulations have been used previously to restrain MTSSL probes dynamics to the experimentally obtained DEER distributions and we explore this avenue to deconvolute the effect of MTSSL probe’s dynamics from the experimental distributional data.

For our reMD simulations we first restrained residue pair 174-466. Since this residue pair is on the extracellular side of the PepT_So_ protein, we chose another pair, 201-364, which had the highest KL divergence on the intracellular side. Hence, our next set of reMD simulations involved two restrained pairs one on each side of the protein. We dubbed these sets of simulations as reMD (1 dist) and reMD (2 dist). While in Figures 4A, B and S18A,B, the distance distributions between the ON-ON atoms of the MTSSL probes show a match with the experimental distribution, the closest-heavy atom distances don’t. In Figure 4A, residue pair 174-466 distribution in the teal violin plot has a single dominant peak with a median value of 3.22 nm, whereas the experimental distribution has two peaks. Moreover, the same peak as seen in unbiased BDDM micelle simulations distribution shown in the yellow violin plot is 4 nm. For comparison, this value is 3.98 nm for our unbiased simulations with a labeled residue pair. In general all three extracellular distances in Figure 4A are lower shifted in reMD simulations. This shift is also seen in Figure 4B for the three extracellular distances. reMD (1 dist) and reMD (2 dist) systems both have a labeled residue pair on the extracellular side which would explain lower shifts for all distance shown on this side of the protein. While the distributions are lower shifted in reMD for the extracellular distances, neither a lower shift not an upward shift is seen in the five intracellular distances in Figures 4A or B. Comparing the residue pair 201-364 in Figures 4A and B, we note that when this residue pair is not restrained (teal violin plot) its mean value is 2.66 nm and when it is restrained this value is 3.54 nm, very close to unbiased simulation value of 3.51 nm.

**Figure 4:**
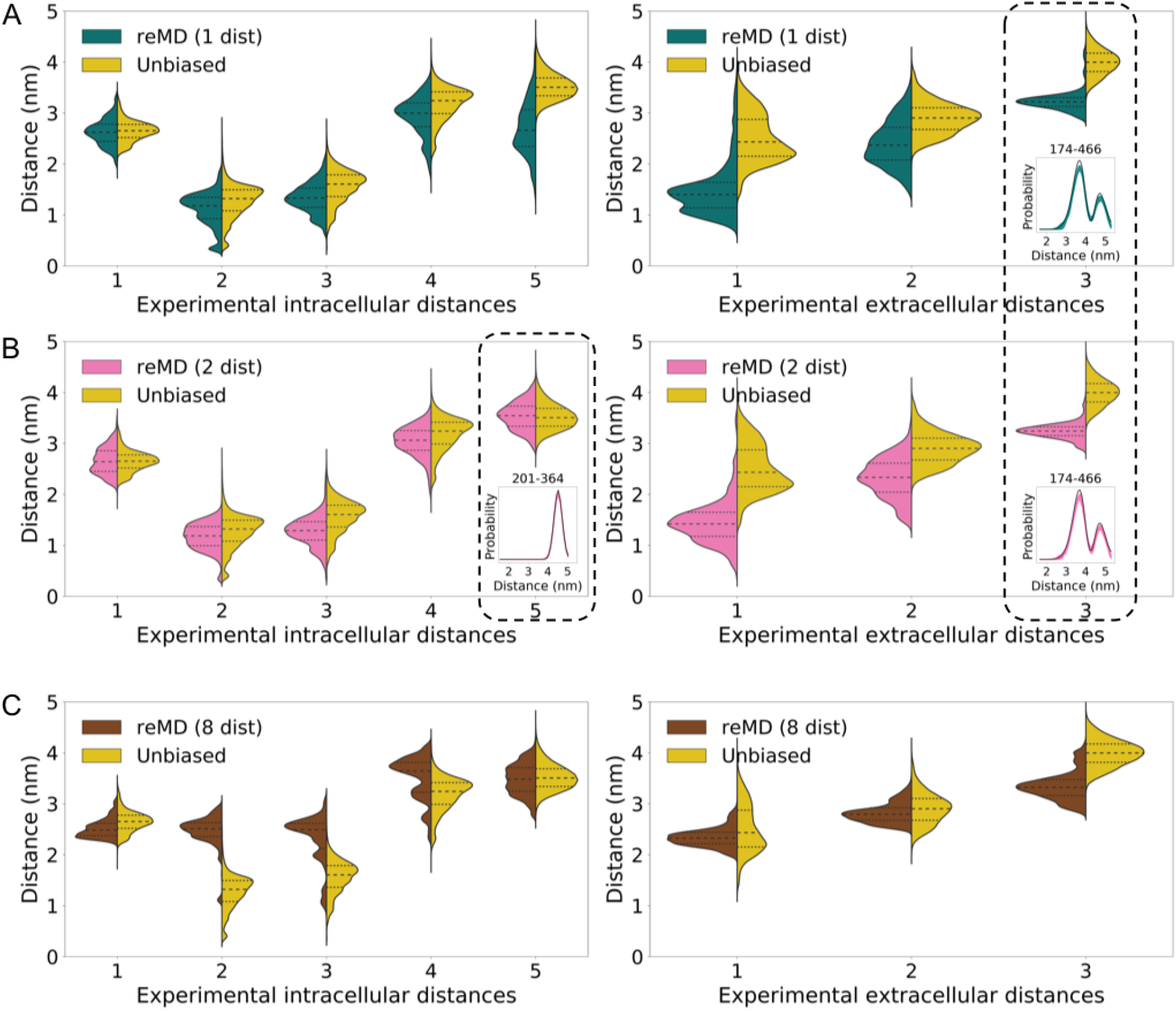
(A) Violin plot shows distance distributions for five intracellular residue-pair distances and three extracellular residue-pair distances as observed from (A) reMD (1 dist) simulations where residue pair 174-466 is restrained, teal violin plots, (B) reMD (2 dist) where residue pairs 174-466 and 201-364 are restrained, pink violin plots, and (C) reMD (8 dist) where all eight residue pairs are restrained, brown violin plots. Yellow violin plots correspond to unbiased simulations of PepT_So_ protein in the micelle. Black dotted outlined residues pairs in (A) and (B) are restrained pairs and probe distances are shown to match with experimental DEER distance distributions.

What happens when we restrain all eight residue pairs in system reMD (8 dist)? Three out of five distances - distance #2, 3, and 4 - on the intracellular side show an upward shift, the median value of the brown violin plots is higher than the median value of the yellow violin plot distributions. Distance #1 and 2 on the extracellular side also are shifted up as compared to systems reMD (1 dist) and reMD (2 dist), although their median values are still lower than those in unbiased simulations.

An upward shift in distance distributions is similar to what we observe in Figure 3 where the ON-ON atom based distances shifted the distribution upwards by ~0.9 nm. However, the origins of these shifts are different. In particular, considering the distance distribution for residue pair 174-466 which is the third distance on the extracellular side, a lower shift in all reMD simulations compared to unbiased simulations without probes (yellow violin plots in Figure 4) and with probes (red violin plots in Figure S18) indicates that reMD simulations alter the backbone dynamics in a way MTSSL probe labeled simulations did not. Vast differences in backbone dihedral angles of the relevant residues in reMD simulations support this observation (Figure S19). These drastic shifts in distance distributions are mirrored in the underlying conformational landscapes (Figure S20). Hence, the bias introduced in reMD simulations via additional energetic terms for force calculations affects the protein structure differently than the modulation caused when MTSSL probes are attached to residues but simulated with unbiased MD simulations.

Similar to our MTSSL-labeled simulations, reMD simulations also suggest that the DEER experiment distance distributions are a convolution of both the spin probe distances as well as the inherent protein dynamics based distances. The impact of spin labels is not straightforward and unbiased simulations are ill-posed to capture their effect completely. Simulations would need to explore the conformational space of each spin label corresponding to every conformational state of the protein. This increases the computational time necessary to capture spin-label dynamics on a protein. With limited computational resources, it is not feasible to perform long-timescale residue pairwise simulations with MTSSL probes. At the same time, while pairwise reMD simulations are also not cheaper, multiple residue pairs can be combined as we demonstrated for our systems reMD (2 dist) and reMD (8 dist). While this may make them computationally tractable, this still does not solve the problem of an unbiased match with MD simulations from long-timescale MD simulations. reMD simulations with multiple restrained residue pairs also raise the unexplored concern that what number of restraints in reMD simulations would be adequate to capture an MD ensemble where all residue-pair distance distributions can correspond to their DEER experiments observables without perturbing the protein’s conformational dynamics.

## Conclusion

This work highlights the necessity for careful interpretation of DEER spectroscopy and MD simulations in membrane protein biophysics. The scarcity of membrane protein biophysical characterization necessitates that we salvage all information available from laboratory experiments and computational simulations. Hence DEER spectroscopy and MD simulations will continue to be important techniques in progressing our understanding of protein dynamics. It is, therefore, imperative to understand how to best compare data obtained from both techniques, not only to show validation of MD simulations with experiments but also to avoid misleading conclusions and to draw predictive conclusions. Previous work has proposed optimization protocols to choose the ideal choice of residue pairs for DEER experiments from already performed MD simulations (*57*, *58*). These protocols can also be used iteratively, performing simulations followed by experiments and then more simulations to update our understanding of a protein’s conformational changes (*59*). Such methods can be used to their full potential once we can decipher the structural characterization of different protein modes identified via multiple peaks in DEER distance distributions. Hence, in this work, we performed a comprehensive study of potential reasons for the discrepancy between DEER experiment distributional data and residue pair distributions from atomistic MD simulations.

We show that the major reason for the difference between experiments and simulation distributions is due to the long length of the MTSSL label and its slow dynamics. The slow dynamics of the flexible MTSSL probes could not be captured in unbiased MD simulations and we examined this using biased simulation methods. While reMD simulations can reconcile experiments and simulations for the restrained residue pairs, reMD yielded significant changes in the protein’s conformational dynamics at the residue-level and globally. It is also not feasible for researchers to perform DEER experiments on all residue pairs of a protein which can be followed by multiple residue pair biased reMD simulations. On the other hand, unbiased MD simulations do not cause any unphysical perturbations in the protein. However, it is computationally expensive to perform long-timescale MD simulations with MTSSL probes. We surmise that when using methods such as *OptimalProbes* (*57*, *59*) it would be sufficient to perform MD simulations with MTSSL probes for the top predicted choices for DEER experiments.

Our computational study of a single pair of spin labels is limited and a thorough examination of the effect of multiple spin labels and different species of spin labels on cytoplasmic and membrane protein is necessary. Our work is also limited in examining the effect of lateral pressure in a bilayer vs micelle environment which deserves examination with different micellar sizes. Our MD simulations are performed at room temperature, Spicher et al. propose performing MD simulations at solvent (mixtures of glycerol and water) freezing temperatures to accurately compare with conformational ensemble explored in DEER experiments (*60*). This is a potential cause for disagreement and is yet to be examined with long-timescale simulations. The simulation temperature will also impact the packing and phase behavior of membranes. Alternatively, using probe molecules such as metal cation based probes which are more rigid (*22*) or employing biophysical experimental methods that do not require any changes to the covalent structure of the target protein that affect the protein’s dynamics and sometimes function could be explored.

## Author Contributions

S.M. and D.S. designed the research. S.M. performed the research, analyzed data, and wrote the manuscript with input from D.S.

## Acknowledgments

D.S. acknowledges support through the New Innovator Award from the Foundation for Food and Agriculture Research and National Center for Supercomputing Applications (NCSA) Fellows Program. S.M. acknowledges support from the University of Illinois Graduate College - Dissertation Completion Fellowship. D.S. and S.M. thank the Blue Waters sustained-petascale computing project, which is supported by the National Science Foundation (Awards OCI-0725070 and ACI-1238993) and the State of Illinois.

