## Supplementary File for "Reconciling Membrane Protein Simulations with Experimental DEER Spectroscopy Data"

### Contents

**Experimental DEER distances** **4**

**MD Simulations** **4**

**Data Analysis** **5**

**References** **33**

6 Residue-pair distance distributions for LeuT simulations in BDDM micelle . 15

7 Residue-pair distance distributions for LeuT simulations in bilayer . . . . . . 16

8 Conformational landscape of $PepT_{So}$ . . . . . . . . . . . . . . . . . . . . . . 17

9 Conformational landscape of LeuT . . . . . . . . . . . . . . . . . . . . . . . 18

10 Comparing mean, median, upper value, and lower value of distance distributions of experimental residue-pair distances and all inter-helix residue-pair distances . . . . . 19
11 Alpha-helical content of TM helices from MD simulations . . . . . 20
12 TM1a alpha-helical content in trajectories started from OF and IF structures of LeuT protein . . . . . 21
13 Alpha-helical content of two short helices in $PepT_{So}$ from MD simulations . . 22
14 Alpha-helical content of intracellular and extracellular loops in LeuT from MD simulations . . . . . 23
15 Ramachandran plots show dihedral angle distributions in micelle and bilayer MD simulations . . . . . 24
16 Distance distributions from MTSSL probes labeled MD simulations for experimental residue-pair distances . . . . . 25
17 Comparing distance distributions between ON-ON atoms and backbone atoms of MTSSL labeled residue pairs. . . . . . 26
18 Distance distributions from restrained-ensemble MD (reMD) simulations for experimental residue-pair distances . . . . . 27
19 Ramachandran plots show dihedral angle distributions in reMD simulations . 28
20 Conformational landscape of reMD $PepT_{So}$ simulations . . . . . 29
21 Residue pair backbone and sidechain distance distributions from MD simulations for experimental distances in $PepT_{So}$ . . . . . 30
22 Residue pair backbone and sidechain distance distributions from MD simulations for experimental distances in LeuT . . . . . 31
23 Multi-Gaussian fits to experimental data . . . . . 32

### Experimental DEER distances

Experimental DEER distances and distance distributions were extracted from previous experiments published in ref. (*1*) and (*2*) using Plot Digitizer Java program.

For $PepT_{So}$, eight DEER distributions are available:

- Five intracellular distances (86-432,141-432,141-438,141-500,201-364)
- Three extracellular distances (47-330,174-401,174-466)

For LeuT protein, we have examined 24 distance distributions because these distributions have data available for Apo system in ref. (*2*).

- 17 intracellular distances (185-271,79-277,184-277,7-86,12-86,12-377,71-193,193-377,12-371,71-89,71-184,71-377,79-377,71-425,71-455,277-425,277-455)
- 7 extracellular distances (309-480,123-240,208-240,37-123,37-208,123-306,208-306)

$PepT_{So}$ experimental distributions were fitted to multiple Gaussian distributions to get an equal-sized-bin distribution for KL divergence calculations and restrained MD simulations. Comparisons shown in Figure S23.

### MD Simulations

##### List of LeuT structural models

2A65 (*3*), 2Q6H (*4*), 2Q72 (*4*), 2QB4 (*4*), 2QEI (*4*), 3F3A (*5*), 3F3C (*5*), 3F3D (*5*), 3F3E (*5*), 3F48 (*5*), 3F4I (*5*), 3F4J (*5*), 3GJD (*6*), 3GWU (*7*), 3GWV (*7*), 3GWW (*7*), 3TT1 (chains A & B) (*8*), 3TT3 (chain A) (*8*), 3USG (*9*), 3USI (chains A & B) (*9*), 3USJ (chains A & B) (*9*), 3USK (chains A, B, C, & D) (*9*), 3USL (*9*), 3USM (*9*), 3USO (chains A & B) (*9*), 3USP (*9*), 5JAE (chains A & B) (*10*), 5JAF (*10*).

### Data Analysis

**Micelle radius**. First, we compute the radius of gyration ($R_g$) of the micelle using *compute_rg* in MDTraj 1.7 (*11*), which is related to the micelle radius ($R$) as, $R = \sqrt{\frac{5}{3}} R_g$ (*12*). This formula hold when the micelle is assumed to be spherical in shape.

**Eccentricity**. The shape of the micelle and the protein-micelle complex is determined using the ratio between moments of inertia $I_1$, $I_2$, and $I_3$ S2. Eccentricity is calculated as $1 - I_{min}/I_{avg}$ (*12*, *13*). The moments are inertia are defined as the eigenvalues of a moment of inertia tensor calculated using *compute_inertia_tensor* in MDTraj 1.7 (*11*).

**Distance distributions**. All inter-residue distance distributions are estimated as the distance between the closest heavy atoms between the two residues unless otherwise mentioned.

**Inter-helix distances** Transmembrane helix ends for proteins are defined based on OPM database (*14*) numbering for 14 helices in $\mathrm{PepT_{So}}$ and 12 in LeuT. We determine inter-helix distances among all helices on intracellular and extracellular side of the proteins. For $\mathrm{PepT_{So}}$ these are $2 \times (14)(13)/2 = 182$ distances and for LeuT these are $2 \times (12)(11)/2 = 132$ distances.

**Kullback-Leibler (KL) divergence**. KL divergence (also called relative entropy) is a measure of how one probability distribution is different from a second, reference probability distribution. KL divergence for two distributions $P$ and $Q$ is 0 if and only if $P$ and $Q$ are exactly equal. For two discrete probability distributions $P$ and $Q$, defined on the same probability space, $X$, the KL divergence of $Q$ from $P$ is defined to be,

$$KL(P|Q) = -\sum_{x \epsilon X} P(x) log(\frac{Q(x)}{P(x)}) \tag{1}$$

KL divergence is an asymmetric measure by definition, and wherever possible we have used this measure both ways to validate our conclusions regarding similarity and difference among probability distribution. We used *scipy.stats.entropy* routine to calculate KL divergence values. Another useful measure of divergence between probability distribution we use is Symmetrised Divergence, which is symmetric and non-negative defined as,

$$\text{Divergence} = KL(P|Q) + KL(Q|P) \tag{2}$$

When calculating frequencies used for the KL divergence we corrected for the presence of frequencies of zero by adding a very small value to the probability distribution.

**Helical content**. The helical content of all TM helices is calculated as defined in the NAMD 2.11 manual (*15*). The python implementation is taken from `https://github.com/amoffett/alpha_helical_content` as used in ref. (*16*). The individual helices in this work are determined based on the OPM database web server (*14*) for $\text{PepT}_{\text{So}}$ PBD 4UVM (*1*) and LeuT PDB 2A65 (*3*). Specifically, TM1 of LeuT refers to residues in TM1a only, which are residues 15 to 25 whereas the helix ends at residue 35.

Table 1: List of MD simulations.

| Complex | Components[#1] | # Trajectories | # Atoms | Equilibration run (ps) | Simulation time (for analysis[#2], $\mu$s) |
|---|---|---|---|---|---|
| LeuT-bilayer | 150 POPE, $Cl^-$ | 72 | 56,707 - 66,784 | 675 | 32.18 |
| LeuT-micelle | 150 BDDM, $Cl^-$ | 72 | 107,521 - 145,589 | 450 | 28.73 |
| $PepT_{So}$-bilayer | 150 POPE, 50 POPG, NaCl | 42 | 66,045 - 74,700 | 675 | 27.3 |
| $PepT_{So}$-micelle | 150 BDDM, NaCl | 42 | 130,506 - 195,681 | 750 | 20.42 |
| $PepT_{So}$-micelle-MTSSL probes | 150 BDDM, NaCl | 42 | 128404 - 181800 | 750 | 18.78 |

#1: All systems contain protein and TIP3P water.
#2: For all trajectories, we eliminate the first 10 ns of the production run from analysis.

Table 2: Geometry of protein-micelle complexes with varied micelle sizes.

| | Complex PepTSo w/ detergents | Micelle Radius (nm) | I1 : I2 : I3 | Eccentricity |
|---|---|---|---|---|
| Micelle | 150 detergents | $4.4 \pm 0.02$ | 1.46 : 1 : 1.24 | $0.23 \pm 0.02$ |
| | 180 detergents | $4.58 \pm 0.03$ | 1.42 : 1 : 1.25 | $0.22 \pm 0.02$ |
| | 200 detergents | $4.68 \pm 0.02$ | 1.53 : 1 : 1.16 | $0.22 \pm 0.02$ |
| Protein+Micelle | 150 detergents | - | 1.26 : 1 : 1.16 | $0.16 \pm 0.02$ |
| | 180 detergents | - | 1.23 : 1 : 1.18 | $0.16 \pm 0.01$ |
| | 200 detergents | - | 1.38 : 1 : 1.13 | $0.17 \pm 0.02$ |

Table 3: List of reMD simulations.

| System | # Trajectories | # Atoms | Equilibration run (ps) | Production run (ns) | Simulation time ($\mu$s) |
|---|---|---|---|---|---|
| reMD (1 dist) | 42 | 9795 | 25 | $\sim$95 | 3.98 |
| reMD (2 dist) | 42 | 11645 | 25 (2 setups required 50 ps) | $\sim$95 | 3.88 |
| reMD (8 dist) | 42 | 19049 | 25 (8 setups required longer) | $\sim$65 | 2.67 |

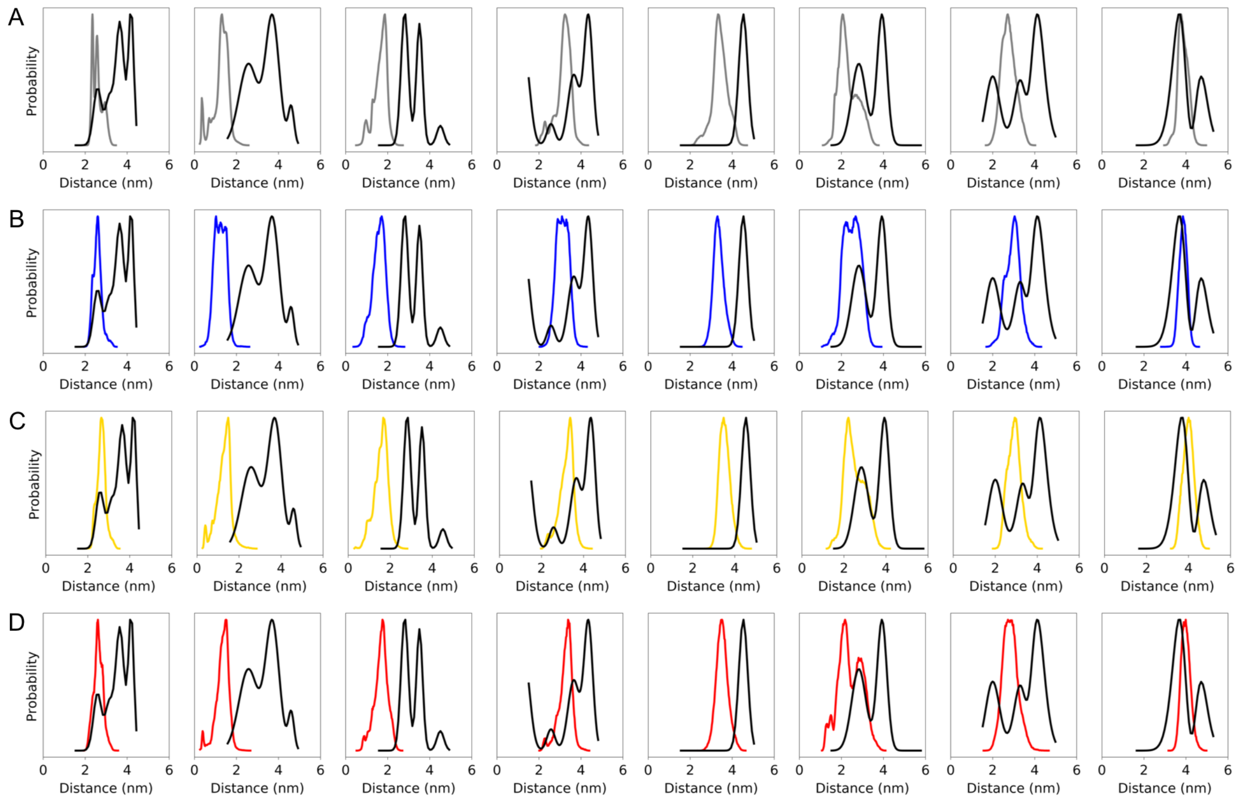

Figure 1: Experimentally characterized residue-pair distance distributions as observed in our MD simulations in (A) POPC bilayer (simulations previously performed (*17*)), (B) POPE/POPG (3:1 ratio) bilayer, (C) BDDM micelle, and (D) BDDM micelle with MTSSL labeled residue pair. Black lines show experimental DEER distance distributions obtained from Fowler et al. as discussed above (*1*). The eight DEER distance distributions shown correspond to distance between residue-pairs 86-432, 141-432, 141-438, 141-500, 201-364, 47-330, 174-401, and 174-466) respectively.

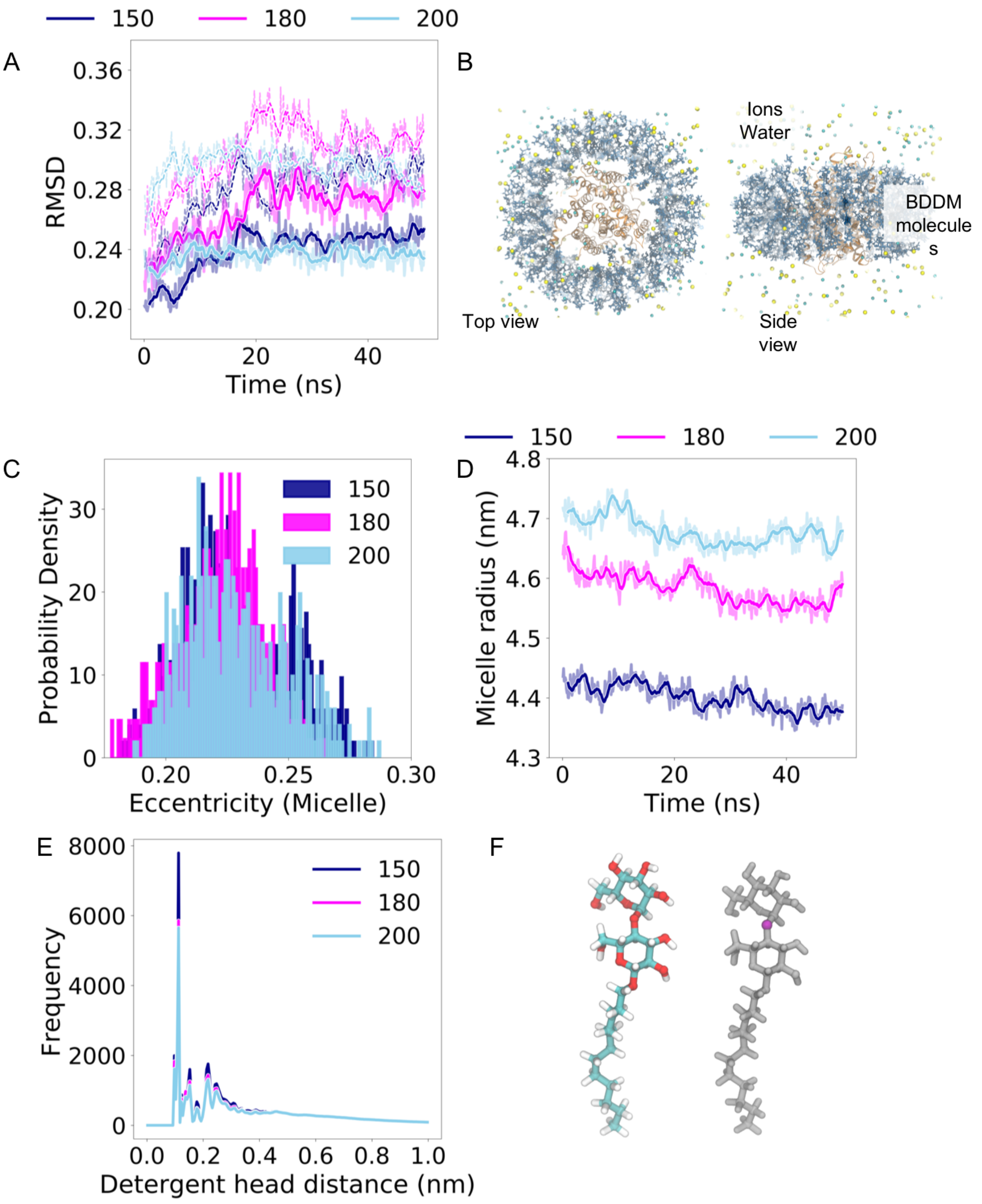

Figure 2: (A) RMSD of protein with respect to the starting frame is shown with time. The dotted lines show RMSD of the full protein while the bold lines show RMSD of the transmembrane region of the protein. Shaded regions show instantaneous values while the lines show a running time average RMSD over a 1 ns time window. (B) An example protein-micelle setup top and side view including BDDM detergent molecules and ions. (C) Probability distribution of micelle eccentricity values. (D) Micelle radius with time is shown. Shaded regions show instantaneous values while the lines show a running time average radius over a 1 ns time window. (E) Radial distribution of distances between BDDM detergent molecule headgroups. Headgroup distances are estimated using the distance among oxygen atoms highlighted in magenta in (F). Colors indicate three micelle sizes, micelle with 150 (blue), 180 (magenta), 200 (sky blue) detergent molecules.

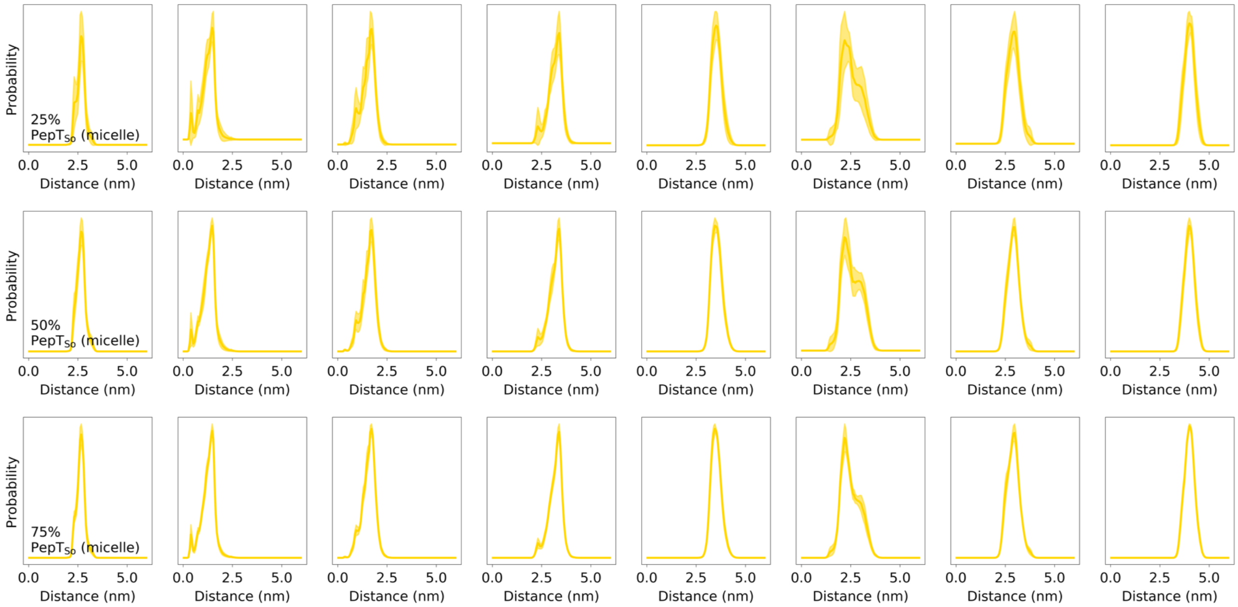

Figure 3: Residue-pair distance distributions for PepT$_{So}$ simulations in BDDM micelle averaged over 25%, 50%, and 75% of the collected trajectories. Filled regions show error bars in the distance distribution as obtained from 10 iterations where a subset of the trajectories is selected randomly. The eight DEER distance distributions shown correspond to distance between residue-pairs 86-432, 141-432, 141-438, 141-500, 201-364, 47-330, 174-401, and 174-466 respectively.

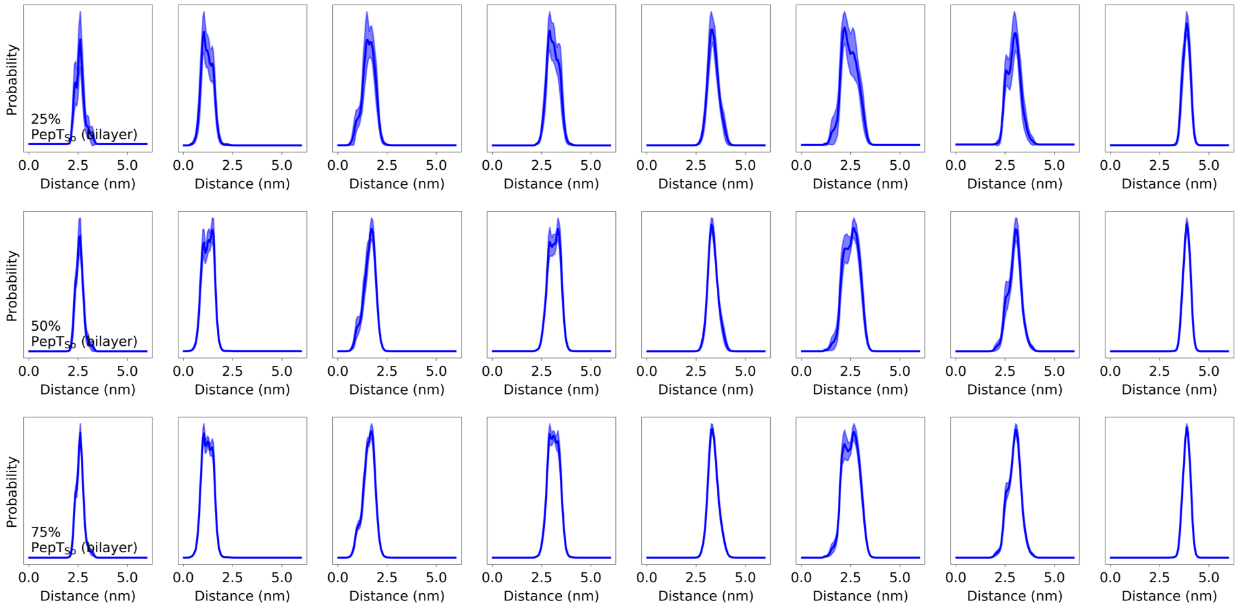

Figure 4: Residue-pair distance distributions for PepT$_{So}$ simulations in bilayer averaged over 25%, 50%, and 75% of the collected trajectories. Filled regions show error bars in the distance distribution as obtained from 10 iterations where a subset of the trajectories is selected randomly. The eight DEER distance distributions shown correspond to distance between residue-pairs 86-432, 141-432, 141-438, 141-500, 201-364, 47-330, 174-401, and 174-466 respectively.

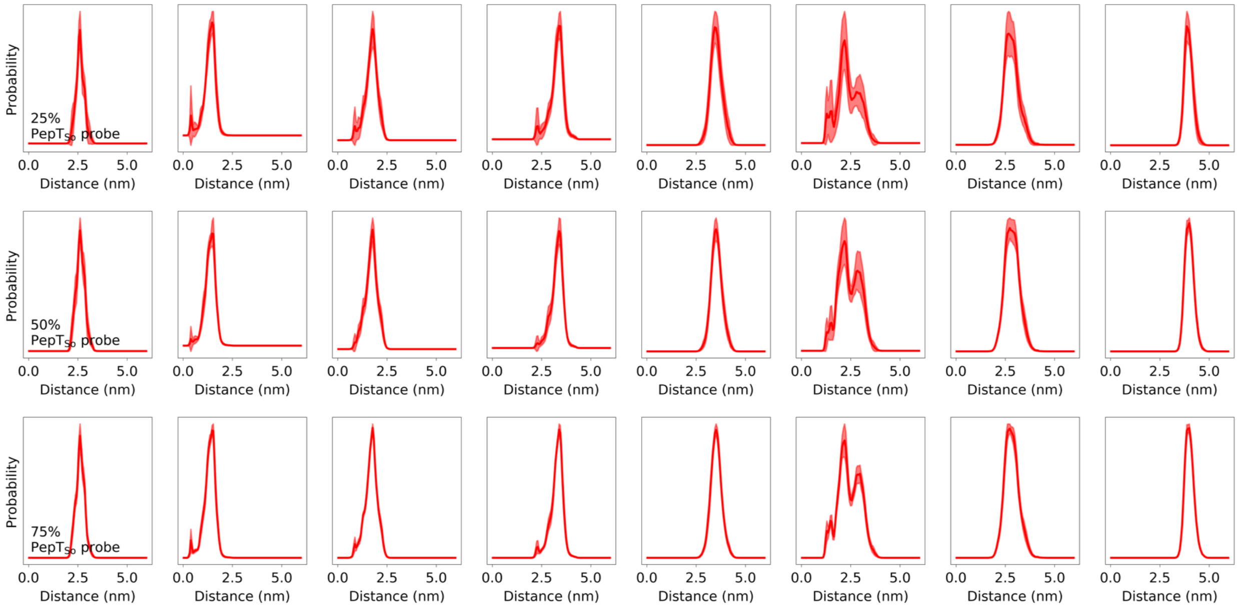

Figure 5: Residue-pair distance distributions for PepT$_{\mathrm{So}}$ simulations in BDDM micelle with a residue pair labeled residue pair averaged over 25%, 50%, and 75% of the collected trajectories. Filled regions show error bars in the distance distribution as obtained from 10 iterations where a subset of the trajectories is selected randomly. The eight DEER distance distributions shown correspond to distance between residue-pairs 86-432, 141-432, 141-438, 141-500, 201-364, 47-330, 174-401, and 174-466 respectively.

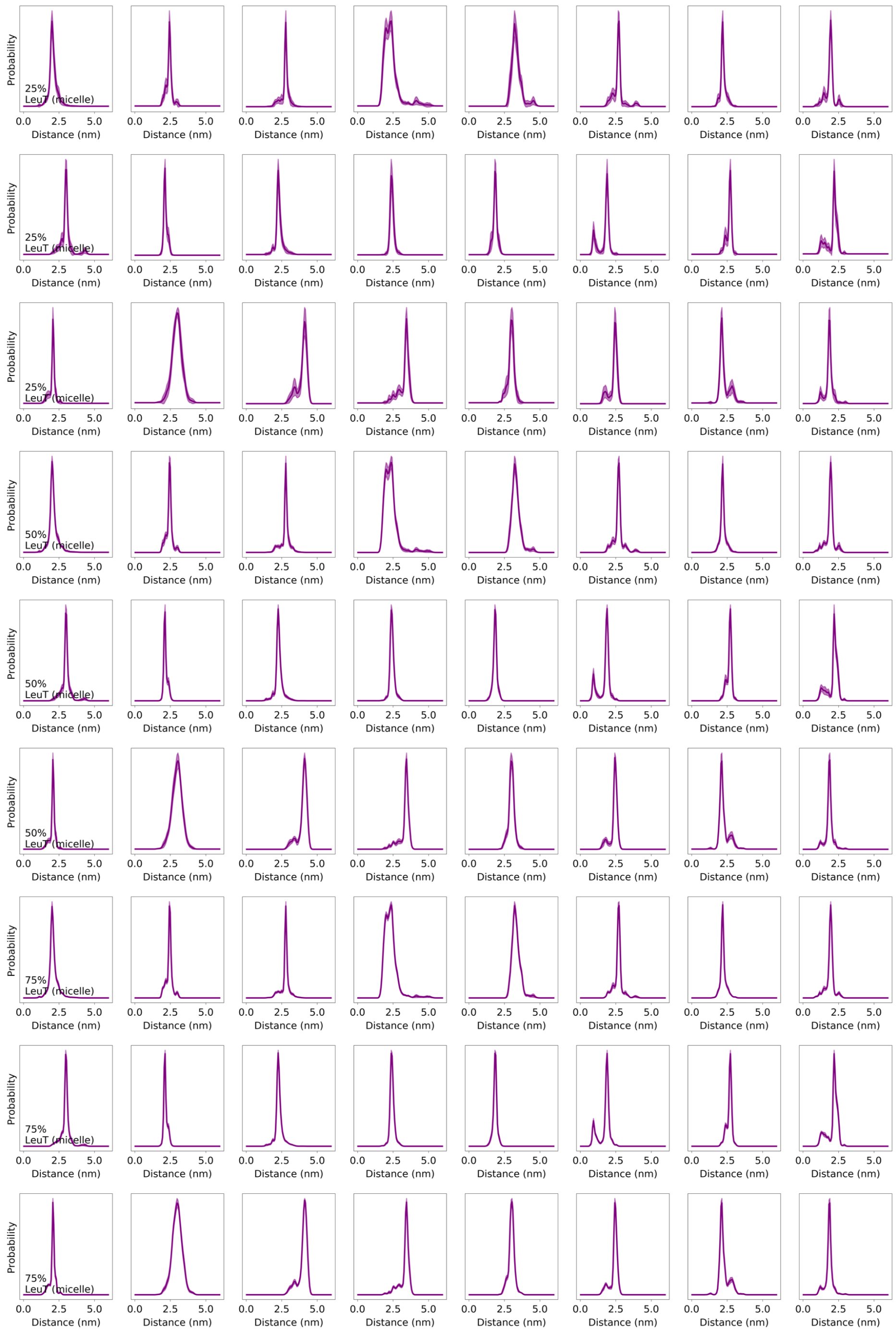

Figure 6: Residue-pair distance distributions for LeuT simulations in BDDM micelle averaged over 25%, 50%, and 75% of the collected trajectories. Filled regions show error bars in the distance distribution as obtained from 10 iterations where a subset of the trajectories is selected randomly. In all 24 DEER distance distributions are shown, where rows 1, 4, and 7 show distances distributions for residue-pairs 185-271, 79-277, 184-277, 7-86, 12-86, 12-377, 71-193, and 193-377; rows 2, 5, and 8 show distance distributions for residue-pairs 12-371, 71-89, 71-184, 71-377, 79-377, 71-425, 71-455, and 277-425; and rows 3, 6, and 9 show distance distributions for residue-pairs 277-455, 309-480, 123-240, 208-240, 37-123, 37-208, 123-306, and 208-306.

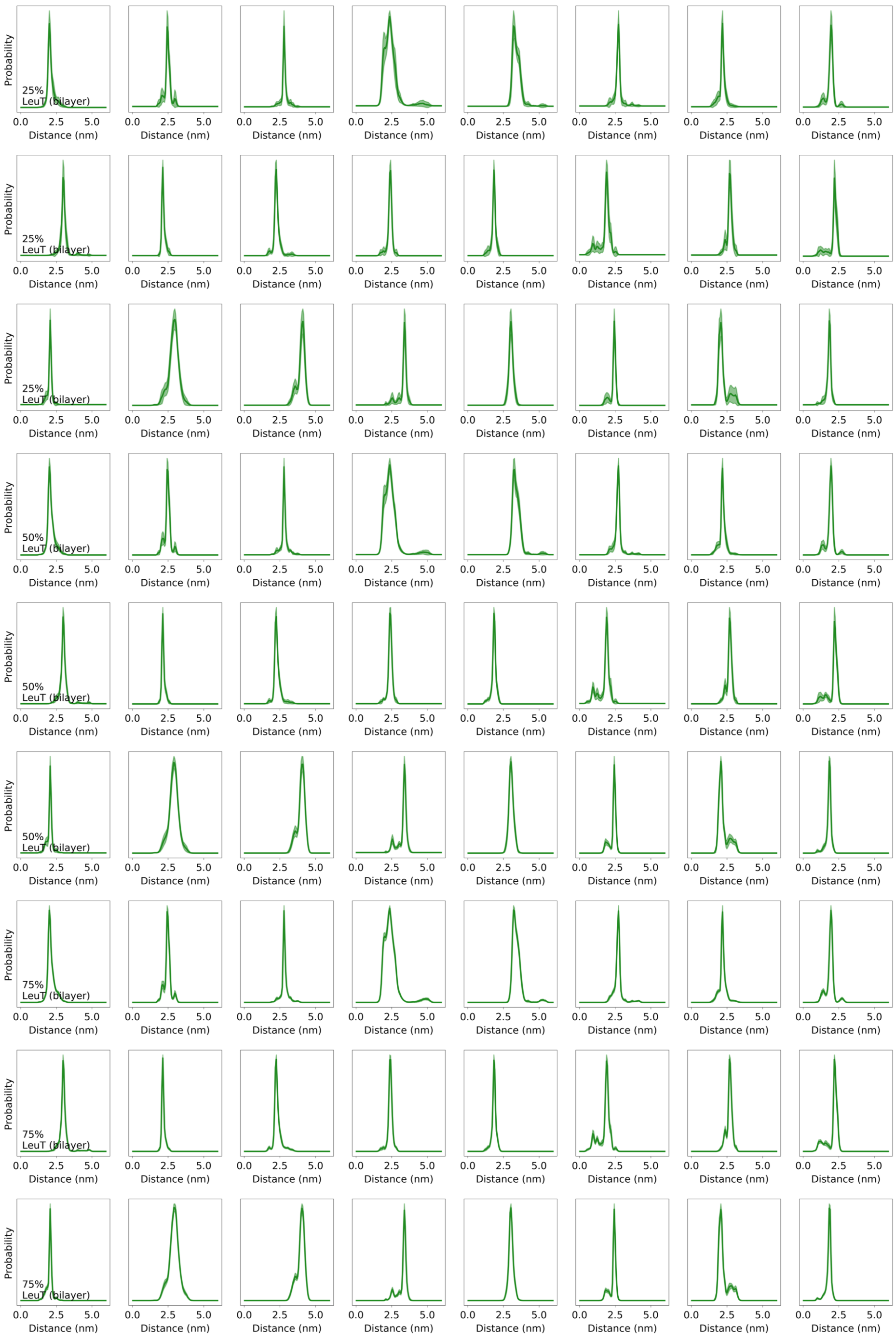

Figure 7: Residue-pair distance distributions for LeuT simulations in bilayer averaged over 25%, 50%, and 75% of the collected trajectories. Filled regions show error bars in the distance distribution as obtained from 10 iterations where a subset of the trajectories is selected randomly. In all 24 DEER distance distributions are shown, where rows 1, 4, and 7 show distances distributions for residue-pairs 185-271, 79-277, 184-277, 7-86, 12-86, 12-377, 71-193, and 193-377; rows 2, 5, and 8 show distance distributions for residue-pairs 12-371, 71-89, 71-184, 71-377, 79-377, 71-425, 71-455, and 277-425; and rows 3, 6, and 9 show distance distributions for residue-pairs 277-455, 309-480, 123-240, 208-240, 37-123, 37-208, 123-306, and 208-306.

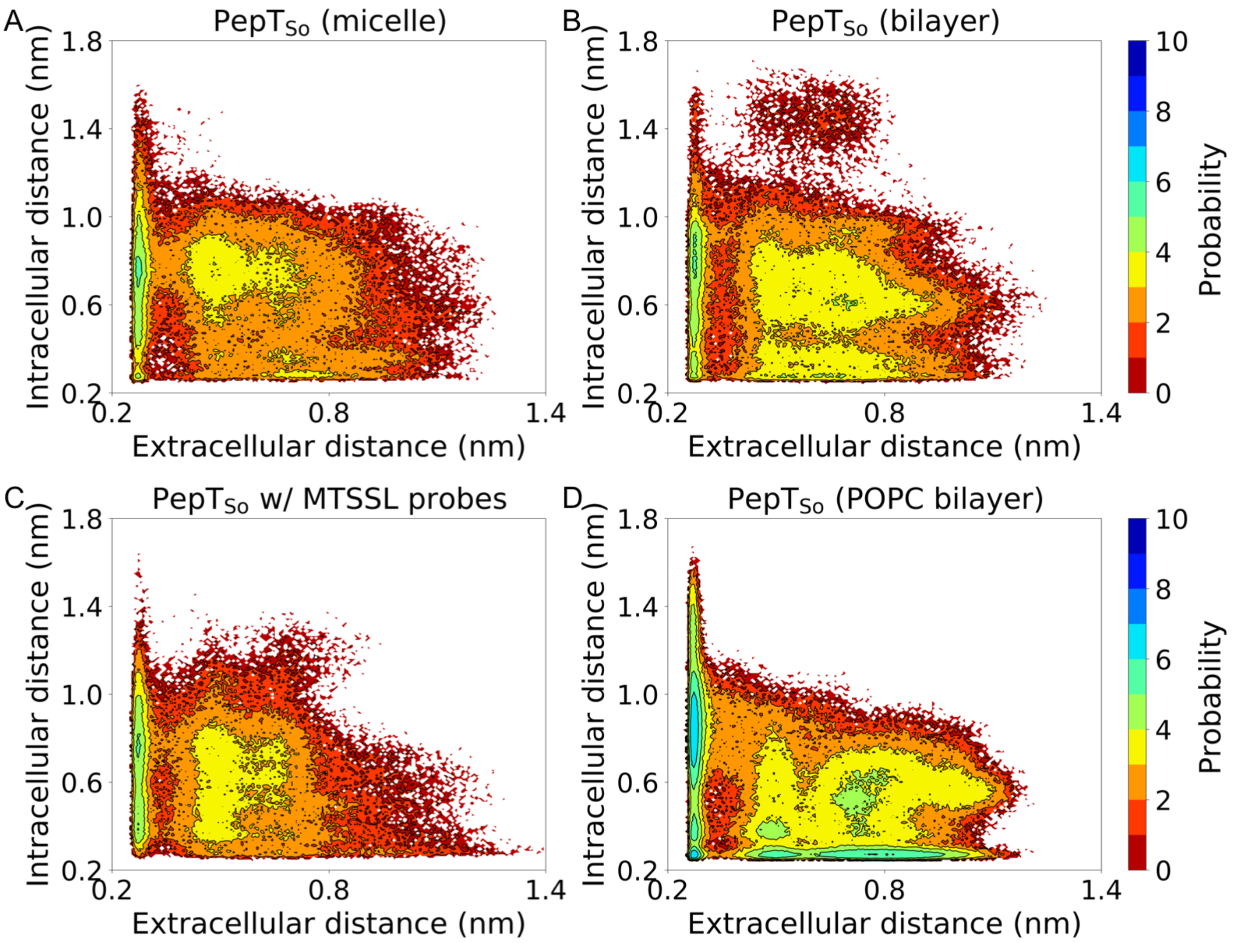

Figure 8: The conformational landscapes of $PepT_{So}$ protein are generated by projecting all simulation data on the chosen extracellular and intracellular side distances measured between Arg32-Asp316 and Ser131-Tyr431, respectively. (A) Conformational landscape for $PepT_{So}$ MD simulations in BDDM micelle. (B) Conformational landscape for $PepT_{So}$ MD simulations in POPE/POPG (3:1 ratio) bilayer. (C) Conformational landscape for $PepT_{So}$ MD simulations in BDDM micelle with an MTSSL labeled residue pair. (D) Conformational landscape from our previous simulations in a POPC bilayer and using an AMBER FF14SB force field (*17*).

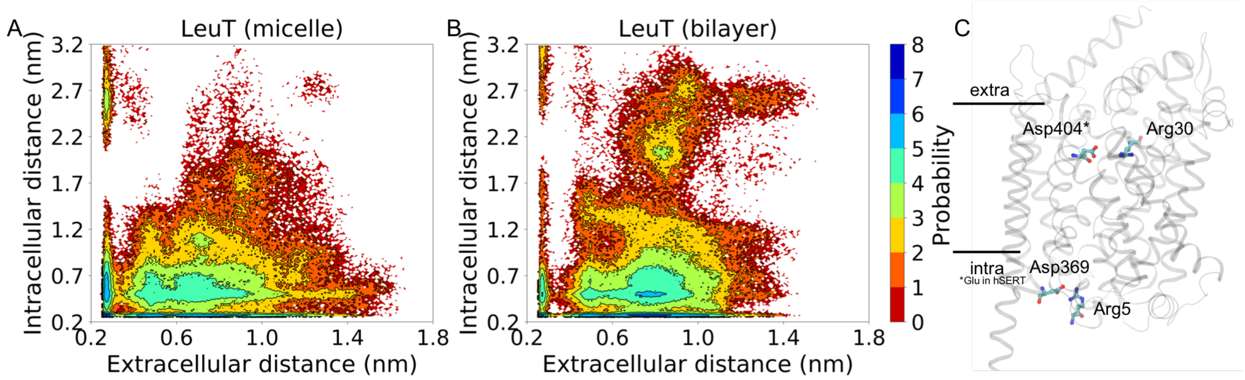

Figure 9: The conformational landscapes of LeuT protein are generated by projecting all simulation data on the chosen extracellular and intracellular side distances measured between Arg30-Asp404 and Arg5-Asp369, respectively. (A) Conformational landscape for LeuT MD simulations in BDDM micelle. (B) Conformational landscape for LeuT MD simulations in a bilayer. (C) Gating residues used to determine extracellular and intracellular distances are shown on a cartoon representation of a three-dimensional LeuT structure.

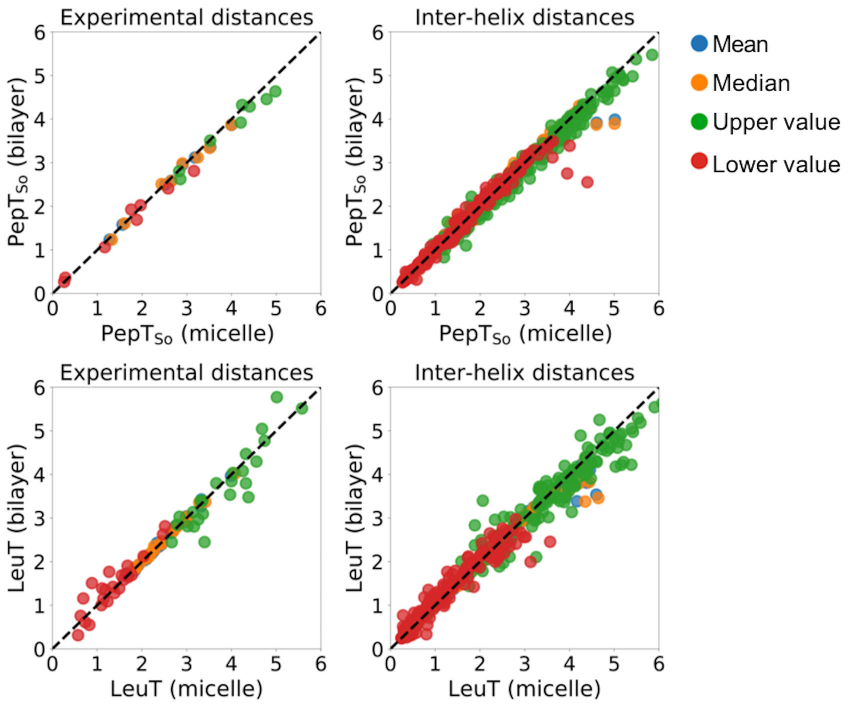

Figure 10: Comparing mean (blue), median (orange), upper value (green), and lower value (red) of distance distributions of experimental residue-pair distances and all inter-helix residue-pair distances. Markers below the black dotted line indicate larger values observed in micelle environment. Markers above the black dotted line indicate larger values observed in bilayer environment. Markers along the black dotted line indicate similar observations in micelle and bilayer simulations.

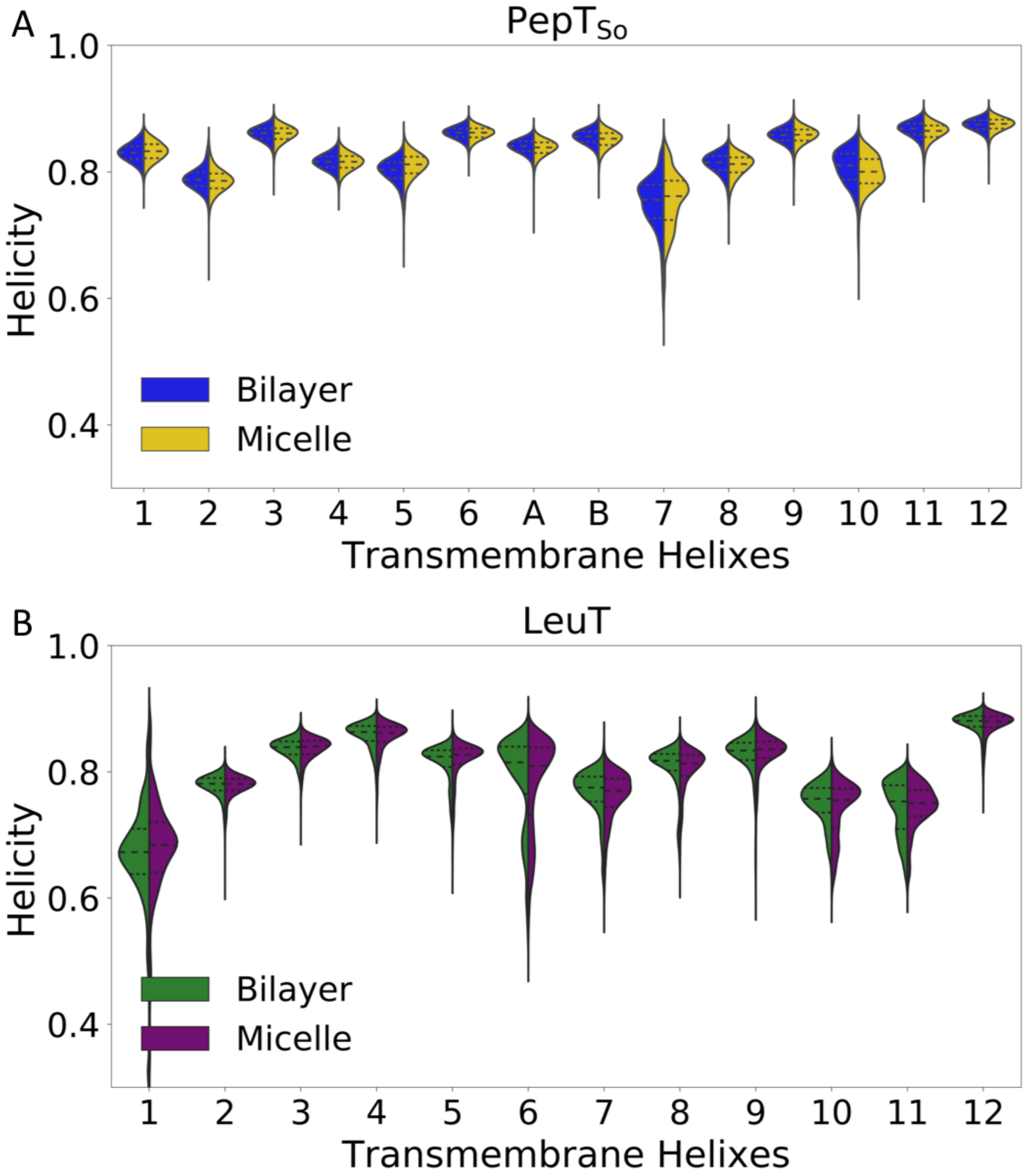

Figure 11: (A) Violin plot shows alpha-helical content for 14 TM helices as observed from MD simulations of PepT$_{So}$ protein in micelle (yellow, right) and bilayer (blue, left). (B) Violin plot shows alpha-helical content for 12 TM helices as observed from MD simulations of LeuT protein in micelle (purple, right) and bilayer (green, left).

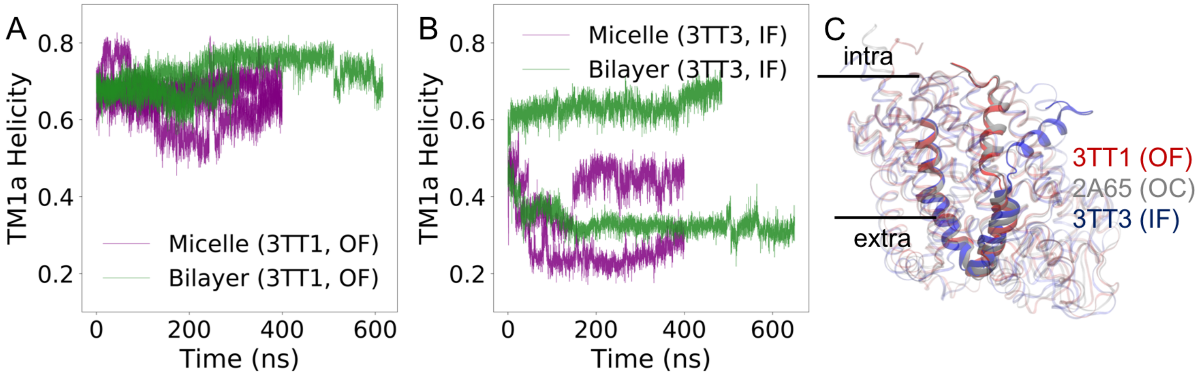

Figure 12: (A) TM1a alpha-helical content of trajectories started from OF structure of LeuT in micelle (purple) and bilayer (green). (B) TM1a alpha-helical content of trajectories started from the IF structure of LeuT in micelle (purple) and bilayer (green). (C) Superposed structures of LeuT's OF, OC, and IF structures.

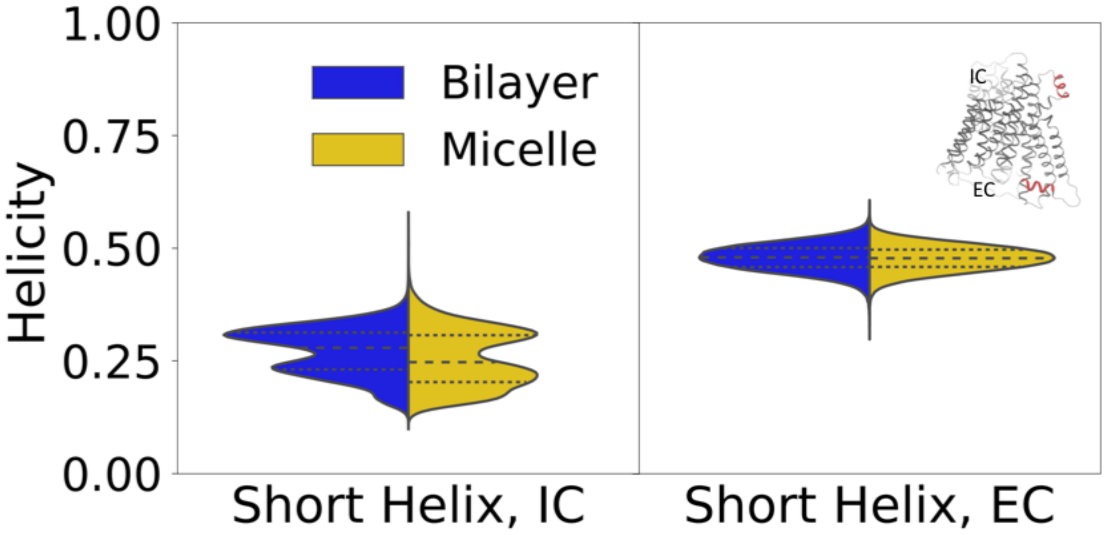

Figure 13: Violin plot shows alpha-helical content of a short helix on the intracellular (IC) side and another of the extracellular (EC) side of $PepT_{So}$ protein in micelle (yellow, right) and bilayer (blue, left). Inset shows two short helices in red on the $PepT_{So}$ protein structure in grey.

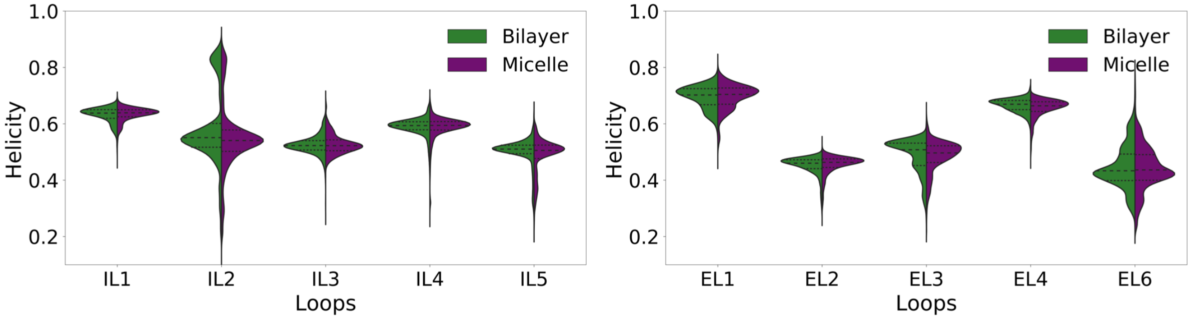

Figure 14: Violin plot shows alpha-helical content of intracellular loops (ILs) and extracellular loops (ELs) in LeuT protein in micelle (purple, right) and bilayer (green, left). Loop EL5 is only four residues long and too short to determine its alpha-helical content.

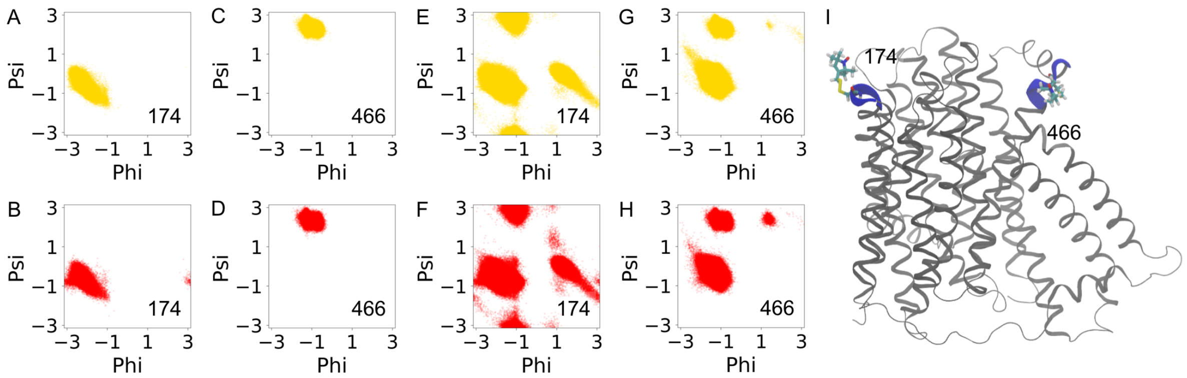

Figure 15: (A-D) Ramachandran plots for residues 174 and 466. Yellow and red colors indicate residue dihedral angle distribution in micelle and bilayer MD simulations, respectively. (E-F) Ramachandran plots for regions surrounding residues 174 and 466. (I) Residues 174 and 466 are shown on a cartoon representation of a three-dimensional $PepT_{So}$ structure.

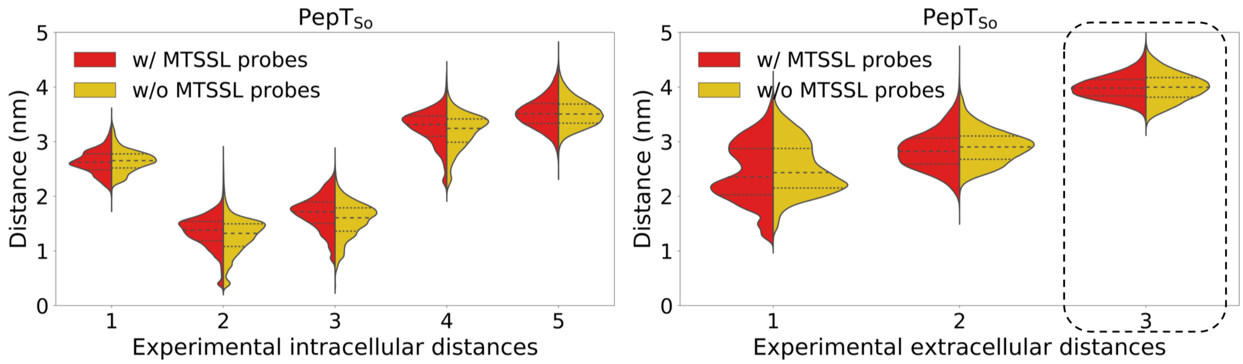

Figure 16: (A) Violin plot shows distance distributions for 5 intracellular residue-pair distances and 3 extracellular residue-pair distances measured by Fowler et al. as observed from MD simulations of PepT$_{So}$ protein in micelle without MTSSL probes (yellow, right) and with an MTSSL probe labeled residue pair (red, left). The black dotted outlined residue pair is the labeled residue pair.

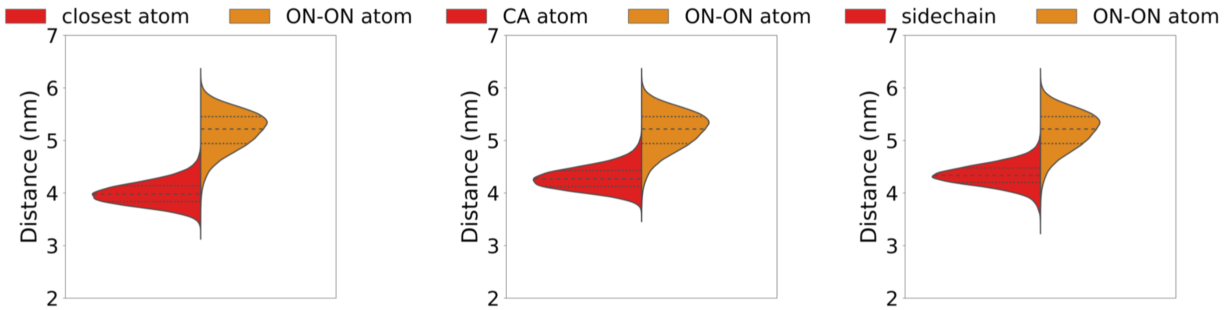

Figure 17: Violin plots compare distance distributions for simulations with MTSSL probes as measured between the ON atom with the closest heavy atom, $C_\alpha$ atom, and the closest sidechain atom of the labeled residues. ON-ON atom distance distributions are shown in orange and the backbone atom distance distributions are shown in red.

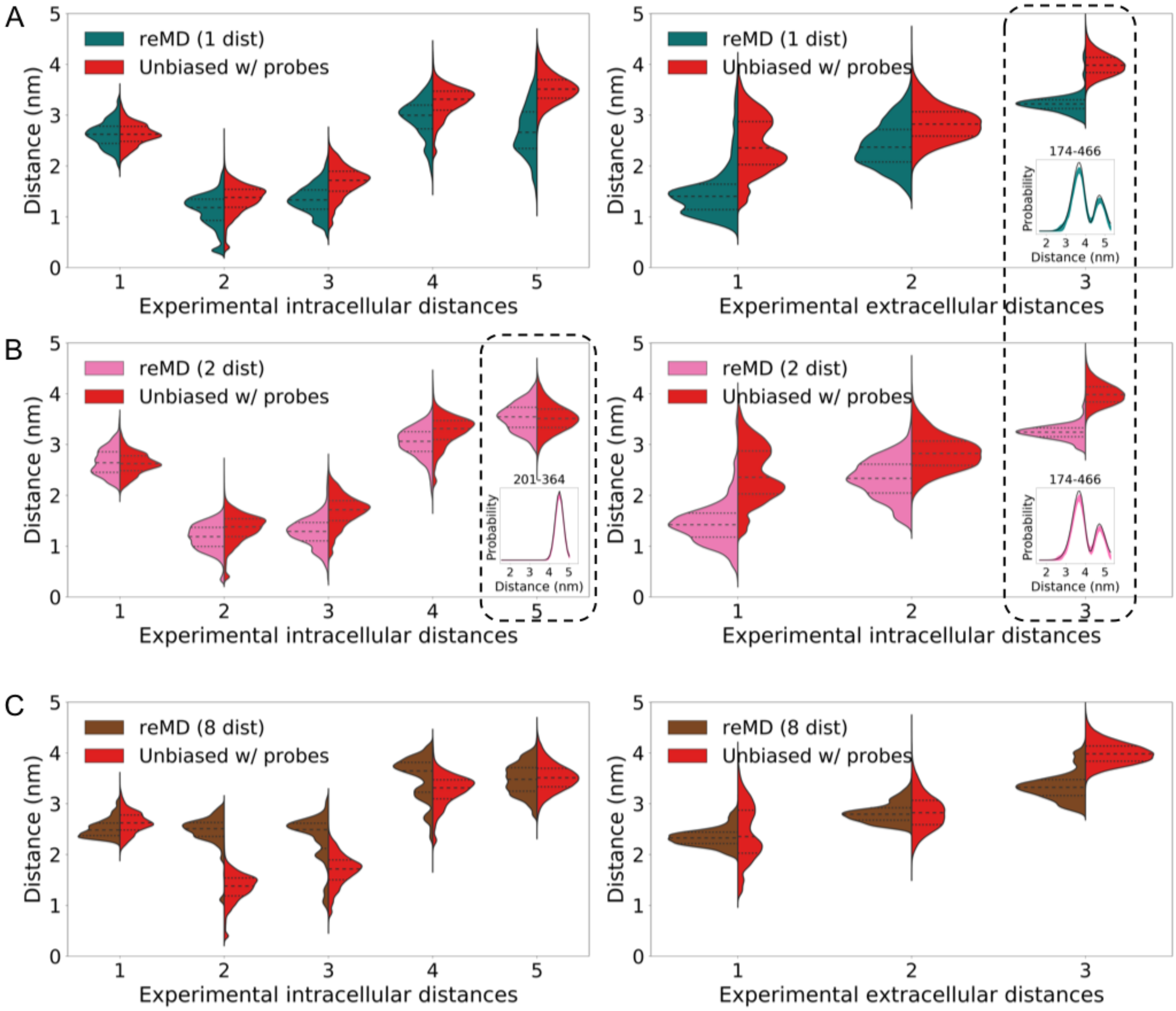

Figure 18: (A) Violin plot shows distance distributions for 5 intracellular residue-pair distances and 3 extracellular residue-pair distances as observed from (A) reMD (1 dist) simulations where residue pair 174-466 is restrained, teal violin plots, (B) reMD (2 dist) where residue pairs 174-466 and 201-364 are restrained, pink violin plots, and (C) reMD (8 dist) where all 8 residue pairs are restrained, brown violin plots. Yellow violin plots correspond to unbiased simulations of $PepT_{So}$ protein in micelle with MTSSL molecules on residues 174 and 466. Black dotted outlined residues pairs in (A) and (B) are restrained pairs and probe distances are shown to match with experimental DEER distance distributions.

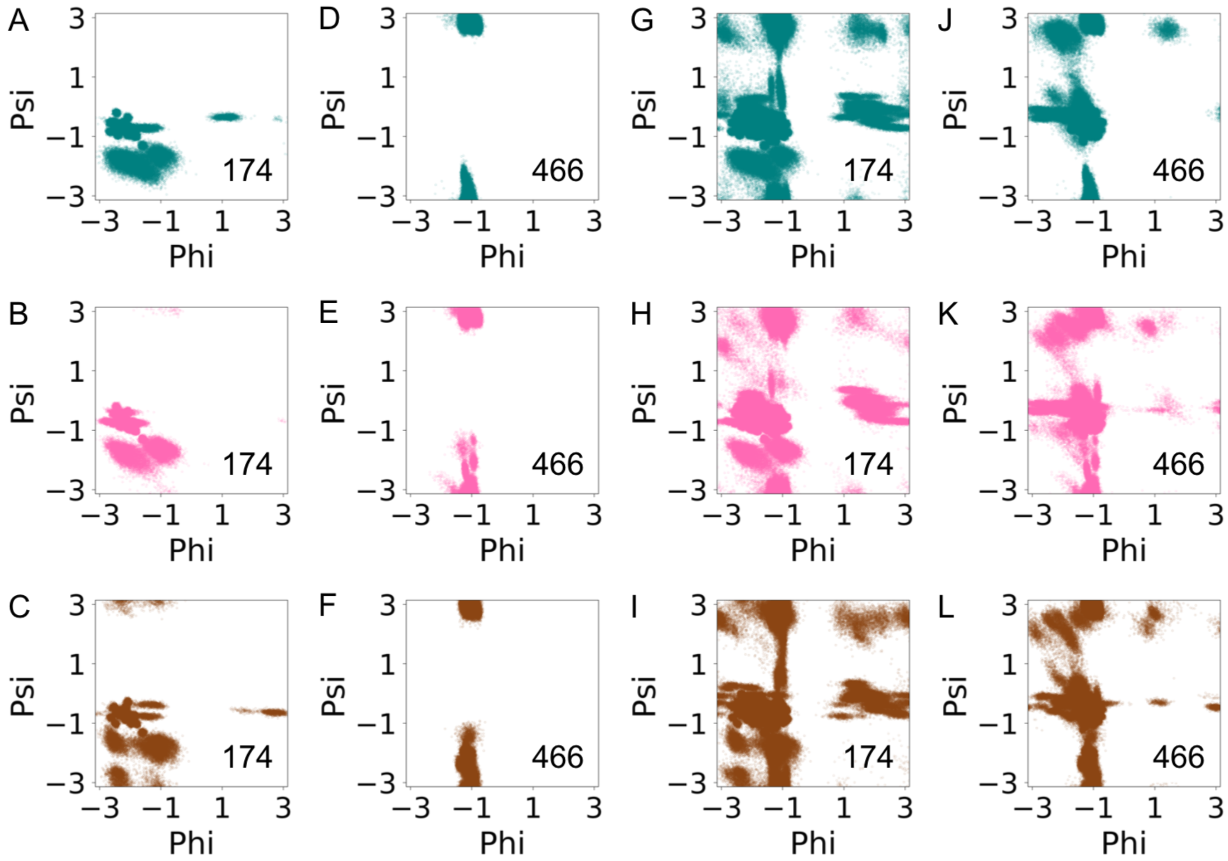

Figure 19: (A-F) Ramachandran plots for residues 174 and 466. Teal, pink, and brown colors indicate residue dihedral angle distribution in reMD (1 dist), reMD (2 dist), and reMD (8 dist) MD simulations, respectively. (G-L) Ramachandran plots for regions surrounding residues 174 and 466.

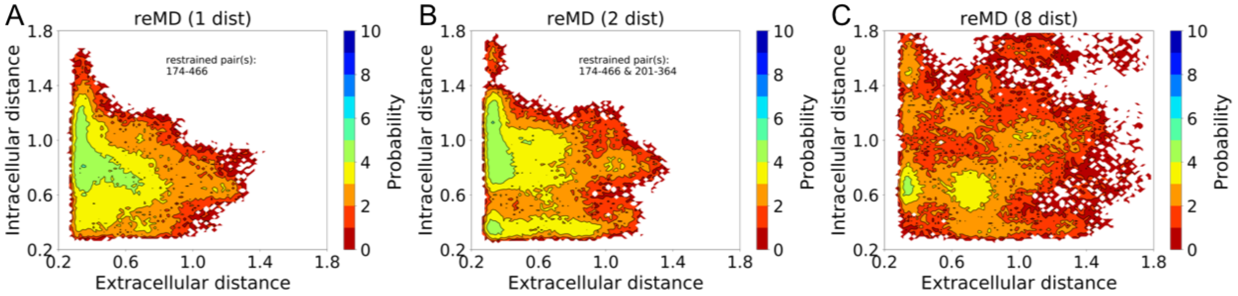

Figure 20: Conformational landscape for $PepT_{So}$ (A) reMD (1 dist), (B) reMD (2 dist), and (C) reMD (8 dist) simulations. The conformational landscapes are generated using the same residue pairs as in Figure S8.

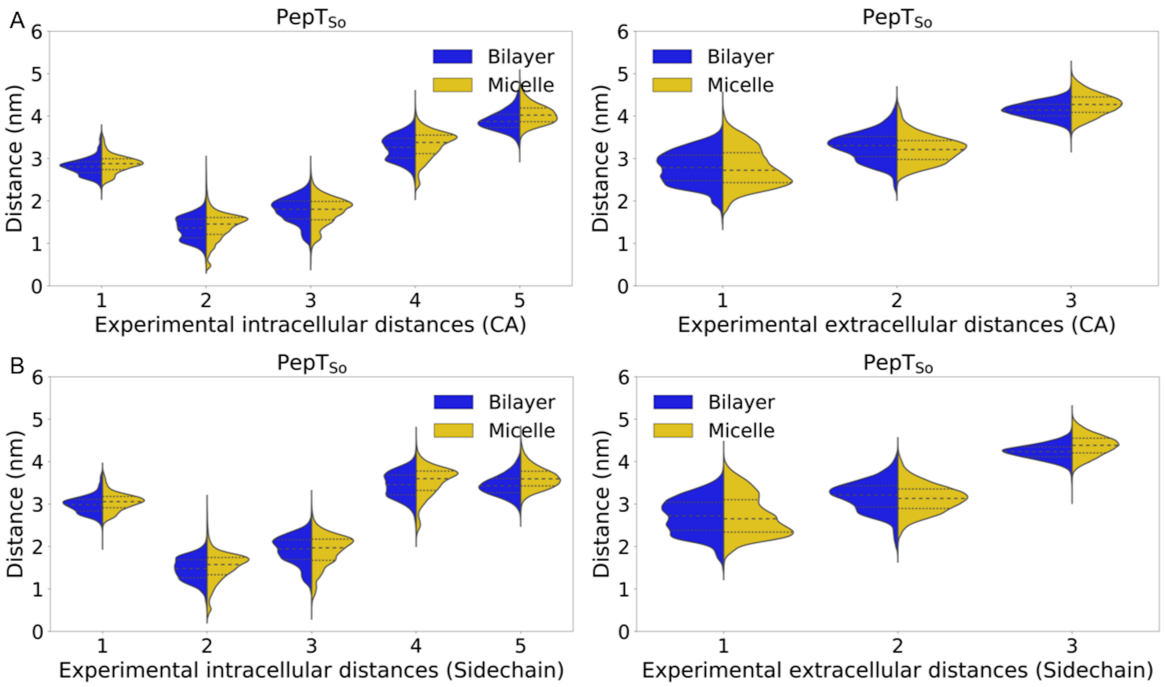

Figure 21: Violin plot shows distance distributions for 5 intracellular residue-pair distances and 3 extracellular residue-pair distances measured by Fowler et al. as observed from MD simulations of PepT$_{\mathrm{So}}$ protein in micelle (yellow, right) and bilayer (blue, left) (*1*). (A) Residue pair backbone distances as measured between $C_\alpha$ atom of residues. (B) Residue pair sidechain distances i.e. closest distance between any two non-hydrogen atoms in residue sidechains.

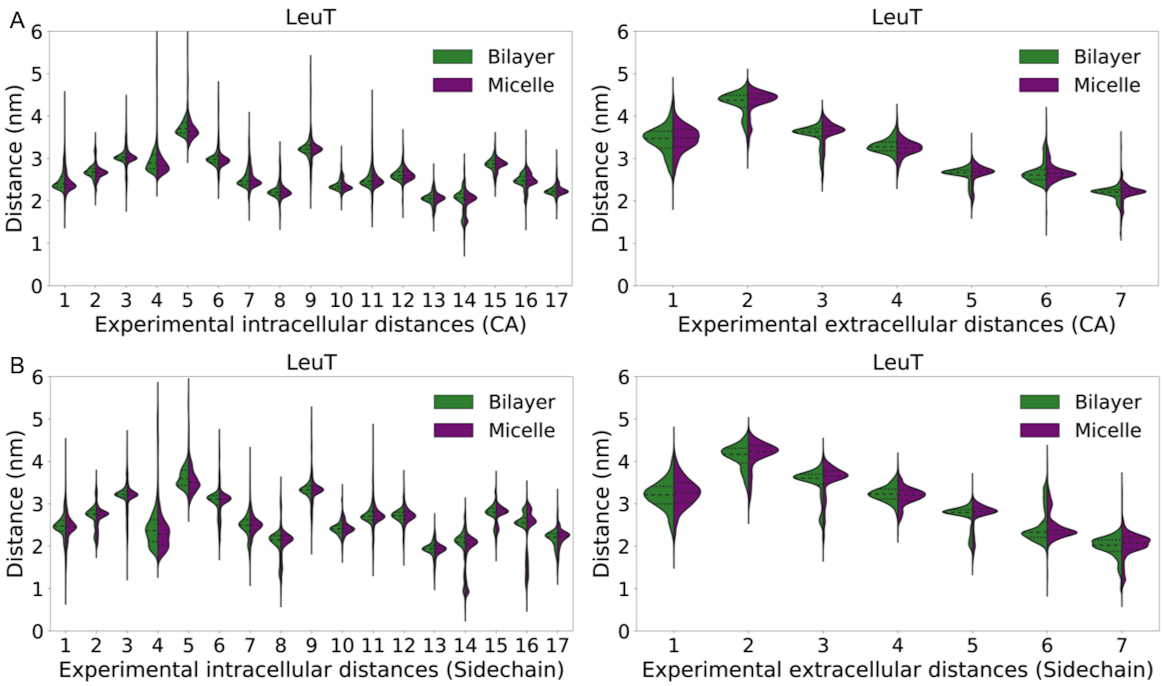

Figure 22: Violin plot shows distance distributions for 17 intracellular residue-pair distances and 7 extracellular residue-pair distances measured by Kazmier et al. as observed from MD simulations of LeuT protein in micelle (purple, right) and bilayer (green, left) (*2*). (A) Residue pair backbone distances as measured between $C_\alpha$ atom of residues. (B) Residue pair sidechain distances i.e. closest distance between any two non-hydrogen atoms in residue sidechains.

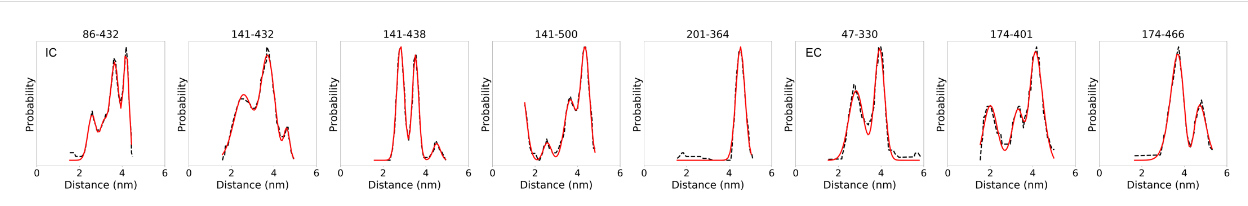

Figure 23: Black dotted lines indicate experimental distributions obtained by tracking data from Fowler et al. and red lines indicate multiple Gaussian fitted to the experimental traces (*1*).
